# FimV, ParC and ParP coordinate polar assembly of the chemosensory arrays in *Pseudomonas putida*

**DOI:** 10.1101/2025.10.27.684771

**Authors:** Marta Pulido-Sánchez, María Zapata-Cruz, Antonio Leal-Morales, Aroa López-Sánchez, Fernando Govantes

**Author notes:** Corresponding author: Fernando Govantes. **Contact:** Marta Pulido-Sánchez, María Zapata-Cruz, Antonio Leal-Morales, Aroa López-Sánchez. Department of Biology, University of Marburg, Marburg, Germany (Marta Pulido-Sánchez).

## Abstract

Motile bacteria rely on chemotaxis to promote directional motion in response to specific environmental signals. The bacterial chemotaxis machinery is typically arranged in highly ordered arrays containing multiple chemoreceptors and signal transduction proteins. *Pseudomonas putida* is a polarly flagellated soil bacterium displaying chemotaxis towards numerous organic compounds. Here we demonstrate the involvement of the polar landmark proteins FimV, ParC and ParP in the assembly of the flagella-associated chemosensory arrays at the cell poles in *P. putida*. Our results show that FimV, ParC and ParP are sequentially recruited to the new cell pole during the cell cycle. FimV is located at the new cell pole concurrently with cell division, and is further required for ParC recruitment. FimV and ParC subsequently stimulate polar localization of ParP. Bacterial two-hybrid assays and structural modeling suggest the involvement of direct protein-protein interactions between the three proteins. We also show that ParC and ParP are decisive to the assembly of fully formed polar chemosensory arrays, by recruiting chemosensory proteins to the cell pole, promoting the stable association of the histidine kinase CheA with the cell pole and enabling the assembly of the functional array structure. Polar targeting of both flagella and chemosensory arrays to the new pole in the timespan of a cell cycle is a means to secure correct segregation of the motility apparatus to both daughter cells during cell division.

**HIGHLIGHTS:**

- FimV, ParC and ParP promote chemosensory array assembly at the new cell pole
- FimV is essential to the normal intracellular distribution of ParC and ParP
- ParC and ParP recruit and stabilize the array components at the cell poles
- ParP and CheA promote polar clustering of the chemoreceptors
- Assembly in the timespan of the cell cycle endows chemotaxis to both daughter cells

**GRAPHICAL ABSTRACT:** 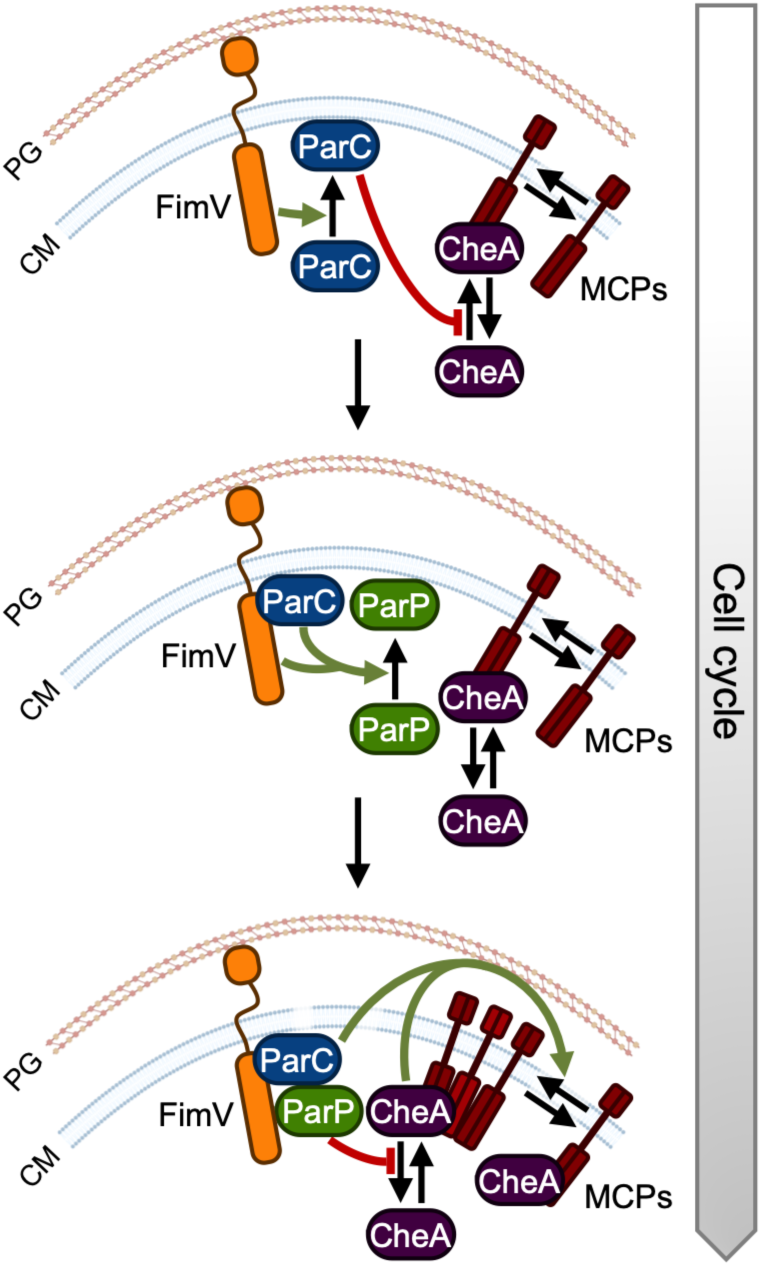

## INTRODUCTION

Motility is a key evolutionary trait that allows bacteria to explore their surroundings **(Colin *et al*., 2021; Miyata *et al*., 2020)**. In many bacterial species, motion is driven by one or several flagella, semi-rigid helical organelles that emerge from the cell body and rotate to propel them in aqueous or semi-solid media **(Altegoer *et al*., 2014)**. Flagellar rotation enables cells to swim in straight runs interrupted by random reorientation events. In response to chemical gradients, bacteria can bias the direction of flagellar rotation to promote efficient navigation towards more favorable environmental conditions. This behavior, known as bacterial chemotaxis, is orchestrated by a conserved set of chemosensors, also known as methyl-accepting chemotaxis proteins (MCPs), and signal transduction proteins assembled in highly organized macromolecular structures known as chemosensory signalling arrays **(Briegel *et al*., 2014; Wadhams and Armitage, 2004; Yang and Briegel, 2020)**.

A typical chemotaxis unit is composed of a set of transmembrane chemoreceptors organized as trimers of dimers that interact in the cytoplasm with the adaptor protein CheW and the histidine kinase CheA **(Alexander *et al*., 2010; Sourjik and Wingreen, 2012)**. Clustering of the core chemotaxis units into hexagonal lattice-like arrays is essential for chemotactic activity, boosting signal sensitivity and integration between different chemosensors **(Wadhams and Armitage, 2004; Sourjik and Wingreen, 2012)**. Similarly structured cytoplasmic arrays associated to soluble chemosensors have also been observed **(Thompson *et al*., 2006; Briegel *et al*., 2014)**. Array assembly as first described in *Escherichia coli* is a stochastic process in which individual chemoreceptor proteins randomly insert into the cell membrane and diffuse to eventually join pre-existing clusters or nucleate the formation of new ones **(Greenfield *et al*., 2009; Shiomi *et al*., 2006; Thiem and Sourjik, 2008)**. Many bacterial species localize their chemosensory arrays at the cell poles, although some (e.g., *E. coli*) also bear additional arrays in lateral positions **(Jones and Armitage, 2015)**. Location at the cell poles is likely a means to guarantee segregation to both daughter cells after division **(Jones and Armitage, 2015)**. Multiple mechanisms have been invoked for the polar recruitment and nucleation of the chemosensory components, including stabilization of the structure by the membrane curvature or by cell pole-specific lipids **(Mileykovskaya and Dowhan, 2000; Thiem *et al*., 2007; Hall *et al*., 2012; Strahl *et al*., 2015)**, or interaction with a variety of polar landmark proteins **(reviewed by Jones and Armitage, 2015)**. In the polarly flagellated *Vibrio cholerae* and *Vibrio parahaemolyticus*, chemotaxis arrays localize at the flagellated pole, but their distribution shifts to bipolar prior to cell division. Polar localization and cluster formation at the new cell pole is dependent on the sequential polar recruitment of HubP, a polar organizer protein, ParC, a ParA homologue conserved in many γ-proteobacteria and ParP, a CheW domain-containing protein **(Ringgaard *et al*., 2011; Yamaichi *et al*., 2012; Ringgaard *et al*., 2014).** ParP is in turn required for proper location of diverse components of the chemosensory arrays in *V. parahaemolyticus* and *V. cholerae* **(Ringgaard *et al*., 2014)**. Together, the ParC/ParP system is proposed to recruit the chemotaxis proteins to the cell pole by a diffusion-and-capture mechanism **(Ringgaard *et al*., 2018)**.

*Pseudomonas putida* is a Gram-negative γ-proteobacterium frequently isolated from soils and root-associated microbiomes, often used as model organism for bioremediation of organic pollutants due to its extraordinary metabolic versatility **(De Lorenzo *et al*., 2024)**. During its planktonic lifestyle, *P. putida* displays a tuft of 3-6 flagella at a single cell pole that is controlled by the chemotactic response **(Harwood *et al*., 1989)**. Research in the past decades has identified numerous *P. putida* soil isolates displaying chemotaxis towards a wide range of aromatic pollutants (**Grimm and Harwood, 1997; Harwood *et al*., 1984, 1990; Lacal *et al*., 2011; Luu *et al*., 2013; Sarand *et al*., 2008)** and other organic compounds present in root exudates **(Canarini *et al*., 2019; López-Farfán *et al*., 2019)**.

The *P. putida* KT2440 genome encodes 27 different chemoreceptors, and the function of 14 of them has been elucidated so far **(summarized in Rico-Jiménez *et al*., 2022)**. In this strain, most flagellar genes and the core components of the flagella-dependent chemotaxis signal transduction system (**García-Fontana *et al*., 2013; López-Farfán *et al*., 2019)** are encoded in a near-contiguous single genomic region known as the flagellar cluster **(Leal-Morales *et al*., 2022)**. A three-tier transcriptional cascade involving the transcription factor FleQ and the alternative σ factor FliA regulates the expression of the flagellar and core chemotaxis signal transduction genes **(Leal-Morales *et al*., 2022)**, and multiple chemoreceptor-encoding genes **(Österberg *et al*., 2010; Rodríguez-Herva *et al*., 2010; Marta Pulido-Sánchez, unpublished).** A second chemotaxis-like system, WspABCDEFR, regulates c-di-GMP synthesis and biofilm development in a flagella-independent fashion **(Hickman *et al*., 2005; Nie *et al*., 2022)**. Orthologs of the *P. aeruginosa* Chp-Pil chemotaxis system regulating type IV pili-dependent surface motility **(Whitchurch *et al*., 2004; Bertrand *et al*., 2010)** are also found in the *P. putida* genomes.

Our recent work showed that deletion of the HubP ortholog FimV provokes cells to swim in short, erratic trajectories, reflecting a defect in flagellar function **(Pulido-Sánchez *et al*., 2025)**. We described that FimV is a polar landmark protein present at both cell poles. FimV is recruited to the division site in predivisional cells to remain permanently attached to the new poles after cell division. FimV is essential to polar localization of the MinD/ParA-type ATPase FleN, a regulator of the number of flagella **(Navarrete *et al*., 2019)**, and they both play a role in synchronizing the assembly of a new set of flagella at the non-flagellated cell pole with the cell cycle. In addition, we showed that FimV stabilizes the polar association of the flagellar landmark protein FlhF, preventing the accumulation of flagellar assembly intermediates at non-polar regions **(Pulido-Sánchez *et al*., 2025)**. Although the intracellular distribution of the core flagella-associated chemotaxis machinery in *P. putida* remains uncharacterized, early studies in *P. putida* reported that the Aer chemoreceptors appear at a single cell pole **(Sarand *et al*., 2008)**, and our recent work unveiled that CheA is located at the flagellated pole in wild-type cells **(Schmidt et al., 2026)**, suggesting that the chemosensory arrays share the polar location of the flagella in this organism.

Here we demonstrate the involvement of FimV, ParC and ParP in the spatial distribution of the chemotaxis arrays in *P. putida* during the cell cycle. Our results indicate that the ParC/ParP system is targeted by FimV to the cell pole, being essential to the chemotactic response by promoting chemoreceptor clustering and stabilizing CheA interaction with the arrays. Our findings shed light on the mechanisms that coordinate the inheritance of the chemosensory arrays associated to the flagellar system in polarly flagellated bacteria.

## MATERIALS AND METHODS

### Bacterial strains and growth conditions

Bacterial strains used in this work are listed in **Table S1.** Liquid cultures of *E. coli* and *P. putida* were routinely grown in lysogeny broth (LB) **(Sambrook and Russell, 2000)** with 180 rpm shaking at 37 °C and 30 °C, respectively. Minimal medium was prepared as previously described **(Mandelbaum *et al*., 1993)** containing lactose (25 mM) and ammonium chloride (1000 mg l^-1^) as the sole carbon and nitrogen sources. For solid media, 1.5% American Bacteriological Agar (Condalab) was added prior to autoclaving. When appropriate, antibiotics and other compounds were added at the following concentrations: ampicillin (100 mg l^-1^), carbenicillin (500 mg l^-1^), chloramphenicol (15 mg l^-1^), gentamycin (10 mg l^-1^), rifampicin (20 mg l^-1^), kanamycin (25 mg l^-1^, or 50 mg l^-1^ when using *E. coli* BTH101 as host strain), 3-methylbenzoate (3MB) (3 mM), 5-bromo-4-chloro-3-indoyl-β-D-galactopyranoside (X-gal) (25 mg l^-1^), isopropyl β-D-1-thiogalactopyranoside (IPTG) (1 mM), sodium salicylate (2 mM). All reagents were purchased from Sigma-Aldrich.

### Plasmid and strain construction

Plasmids and oligonucleotides used in this work are listed in **Table S1**. DNA manipulations were conducted following standard protocols **(Sambrook and Russell, 2000)**. Restriction and modification enzymes were used following the manufacturer’s instructions (New England Biolabs). PCR amplifications were performed using Q5^®^ High-Fidelity DNA Polymerase (New England Biolabs) for cloning purposes, and DreamTaq^TM^ DNA Polymerase (Thermo Fischer) to check plasmid constructions and chromosomal manipulations. *E. coli* DH5α was used as host strain for cloning procedures. Cloning steps involving PCR were verified by commercial Sanger sequencing (Stab Vida). Plasmid DNA was transferred to *E. coli* strains by transformation **(Inoue *et al*., 1990)** or electroporation **(Dower *et al*., 1988)**, and to *P. putida* by triparental mating **(Espinosa-Urgel *et al.,* 2000)** or electroporation **(Choi *et al.,* 2006)**. Site-specific integration of miniTn*7*-derivatives at the single *att*Tn*7* chromosomal site in *P. putida* strains was performed and PCR-verified as previously described **(Choi *et al.,* 2005)**. Specific details of plasmid and strain construction are provided as **Supplementary Materials and Methods**.

### Swimming motility assays

Swimming assays were adapted from Parkinson **(Parkinson, 1976)** Fresh colonies were toothpick-inoculated in Tryptone-agar plates (1% Bacto^TM^ Tryptone [Difco], 0.5% NaCl, 0.3% Bacto^TM^ Agar), and incubated at 30 °C for 12 h or 48 h. Medium was supplemented with sodium salicylate when required. Digital images were taken and swimming halos diameter were measured using GIMP v2.10.10 **(The GIMP Development Team, 2019)** and normalized to the wild-type.

### Chemotaxis assays

Quantitative chemical-in-plug chemotaxis assays were performed as previously described **(Parales and Ditty, 2018)** with some modifications. Plugs of 2% agarose prepared in chemotaxis buffer (25 mM Na_2_HPO_4_, 25 mM KH2PO4, 0.05% glycerol, 10 μM EDTA, pH 7.0) containing 0.2% casamino acids as chemoattractant were placed onto the center of 60×15 mm Petri dishes. Plates were poured with 10 ml of chemotaxis buffer containing 0.2% noble agar and allowed to solidify at room temperature for 1 h. Overnight cultures were 250-fold diluted in LB medium and grown to exponential phase (OD_600_ of 0.4-0.5). Cells were harvested by centrifugation (3500 *g*, 10 min, room temperature), washed three times with 1 ml of chemotaxis buffer by gently pipetting to minimize flagella breakage, and adjusted to an OD_600_ of 20. Plates were inoculated by pipetting 2 µl of cell suspensions into the soft agar medium and incubated at 30 °C for 12 h. Digital images and halo measurements were taken as described above. Chemotaxis response indexes were calculated as the distance from the inoculation point to the edge of the motility halo closest to the agarose plug, divided by the distance from the inoculation point to the edge of the halo farthest from the agarose plug.

### SDS-PAGE and Western-Blot analysis

Cell pellets and protein samples were mixed with 2x loading buffer (150 mM Tris-HCl pH 6.8, 20 % glycerol, 4% SDS, 0.1 % bromophenol blue, 10 % β-mercaptoethanol) and boiled at 100 °C for 10 min. Samples were centrifuged at 18000 *g* for 1 min and supernatants were separated on 12.5 % SDS-PAGE gels. Proteins were transferred to nitrocellulose membranes using a Trans-Blot Turbo Transfer System (Bio Rad). Protein transference was evaluated by Ponceau S Staining Solution (Thermo Fischer). Membrane was blocked for 2 h at room temperature in blocking solution (3 % fat-free milk in TBS buffer with 1 % Tween 20). Blocked membrane was incubated overnight at 4 °C with primary anti-GFP antibody (SAB4301138, Sigma-Aldrich, 1:5000 dilution), washed 3 times in TBS with 1% Tween 20 (TBS-T), probed for 2 h at 4 °C with secondary anti-rabbit (31460, Thermo Fischer, 1:10000 dilution) HRP-conjugated antibody, and washed with TBS-T as above. Pierce™ ECL Western Blotting Substrate (Thermo Fischer) was then added to the membranes and the chemiluminescent signal was revealed using a ChemiDoc MP instrument (Bio-Rad) with a exposure time of 800 s. Image Lab software (Bio-Rad) was employed for image analysis.

### Bacterial Adenylate Cyclase Two-Hybrid (BACTH) assays

Protein-protein interactions were analyzed using a bacterial two-hybrid approach as previously described **(Karimova *et al*., 1998).** Briefly, the proteins to be tested were fused to the T18 and T25 fragments of the adenylate cyclase catalytic domain of *Bordetella pertussis*. Electrocompetent *E. coli* BTH101 cells were prepared as described **(Karimova *et al*., 1998; 2000)** and transformed with the corresponding pUT18 and pKNT25/pKT25-derived plasmids by electroporation. Selection LB plates containing ampicillin and kanamycin were incubated at 30°C for 48 h. Three isolated colonies per combination were inoculated into the wells of a 96-well polystyrene plate containing 150 μl LB supplemented with ampicillin, kanamycin and IPTG, and subsequently incubated at 30 °C with 180 rpm shaking for 24 h. Drops containing 10 µl of each culture were spotted on minimal lactose agar plates supplemented with IPTG and X-gal. Plates were incubated at 30°C for 4 days and the resulting spots were documented by digital imaging. To quantify the adenylate cyclase activity reconstructed by the interacting hybrid proteins, 100 μl samples withdrawn from the 96-well plates were permeabilized with sodium dodecyl sulfate and chloroform, and β-galactosidase activity was determined as described **(Miller, 1993)**.

### Tracking of swimming trajectories

Continuous videos of near-surface cell motility were recorded using a Leica DMI4000B inverted microscope equipped with a Leica HCX PL FLUOTAR 20x/0.40 CORR PH1 objective lens and a Leica DFC360 FX CCD camera (pixel size: 0.2012 µm). Overnight LB cultures were 250-fold diluted in the same medium and grown to an OD_600_ of 0.3. Serial dilutions 10^3^ to 10^5^ were prepared and 150 µl samples were transferred to the wells of a Costar^®^ 96-well polystyrene plate (Corning). Brightfield videos of cells swimming on the plane of the well surface were recorded for 1 min, and converted to 4 fps time-lapses using ImageJ (Fiji) v1.53t **(Schneider *et al*., 2012)**. Trajectories of 20 cells of each strain were tracked for 15 s using the Manual Tracking v2.1.1 plugin.

### Confocal microscopy and image analysis

Overnight cultures were 100-fold diluted in LB medium supplemented with 2 mM salicylate when necessary and grown to an OD_600_ of ∼0.5. Cells from 1 ml samples were harvested by centrifugation (5000 *g*, 5 min, room temperature) and washed three times with 1 ml PBS. Flagellar filaments of strains expressing FliC^S267C^ were stained by carefully resuspending cells in PBS supplemented with 5 μg/ml Alexa Fluor^TM^ 488 C5 Maleimide (Molecular Probes) **(Hintsche *et al*., 2017)**. After 5 min incubation in the dark, cells were washed three times for excess dye removal. For membrane staining, cells were resuspended in 100 μL of PBS containing 5 μg/ml FM™ 4-64 (Molecular Probes) and incubated for 5 min in the dark. Drops of 2 μL samples were immediately placed on 1% agarose pads prepared with deionized water to be imaged using an Axio Observer7 confocal microscope (Zeiss) equipped with a CSU-W1 spinning disk module (Yokogawa). Images were taken using a Zeiss α Plan-Apochromat 100x/1.46 Oil DIC M27 objective lens and a Prime 95B CMOS camera (Teledyne Photometrics, pixel size: 0.1099 μm), controlled by the SlideBook 6 software. GFPmut3 and Alexa Fluor^TM^ 488 fluorescence were excited with the 488 nm laser line and collected with emission filter 525/50 nm, setting exposure times at 300 and 100 ms, respectively. YFP fluorescence was excited at 514 nm and collected with a 525/50 nm filter (300 ms exposure time). FM™ 4-64 was excited at 561 nm and fluorescence was collected with a 617/73 filter (1000 ms exposure time). For each image, seven Z-sections were taken with a step size of 0.33 μm.

Microscopy images were processed using ImageJ (Fiji) v1.53t **(Schneider *et al*., 2012)** and analyzed using the MicrobeJ v5.13l plugin **(Ducret *et al*., 2016)** following a previously described pipeline **(Pulido-Sánchez *et al*., 2025)**. Additionally, cell contour axis profiles were recorded with MicrobeJ using default parameters and setting a dilate value of 0.2 µm. **Bioinformatics analyses**

Sequence alignments were carried out using the MUSCLE tool from the EMBL-EBI website (https://www.ebi.ac.uk/jdispatcher/msa/muscle) **(Madeira *et al*., 2024)**, and further analyzed using Jalview 2.11 **(Waterhouse *et al*., 2009)**. Similarity between *P. putida* KT2440 and *V. cholerae* O1 El Tor ParC and ParP proteins was scored using the BLASTp tool from the NCBI website (blast.ncbi.nlm.nih.gov/Blast.cgi) **(Altschul *et al*., 1990).** AlphaFold 3-based protein complex modelling **(Abramson *et al*., 2024)** was performed at the AlphaFold server **(alphafoldserver.com)** using random seeds and analyzed with UCSF ChimeraX (Copyright University of California Regents), version 1.11.

### Statistical analysis

Statistical data treatment was performed using GraphPad Prism software, version 8.3.0. Results are reported as the average and standard deviation of at least three biological replicates, or as the median and 95% confidence interval for the median in foci fluorescence quantification. Significance (p-values) of the differences among strains/samples was evaluated by means of the two-tailed Student’s t-test for unpaired samples not assuming equal variance.

## RESULTS

### ParC and ParP are essential for chemotaxis-mediated motility

Homologs of *parC* and *parP* are found within flagella/chemotaxis operons in numerous polarly flagellated γ-proteobacteria **(Ringgaard *et al*., 2011; Ringgaard *et al*., 2014)**. The *P. putida* KT2440 flagellar cluster ORFS PP_4334 and PP_4333 **(Leal-Morales *et al*., 2022) (Fig. 1A)** exhibit 51% and 38% identity, and 70% and 56% similarity to *V. cholerae* ParC (VC_2061) and ParP (VC_2060), respectively **(Fig. S1)**. To first identify whether these two proteins contribute to chemotaxis-dependent motility of *P. putida*, we engineered derivatives of KT2442 (a spontaneous rifampicin-resistant mutant of the reference strain KT2440, used here as wild-type) bearing complete deletions of *parC* and *parP*. The Δ*parC* and Δ*parP* mutants displayed 52% and 85%-reduced swim halo diameters on soft agar plates, respectively, compared to the wild-type strain **(Fig. 1B and C)**. The distribution of flagellar filament tufts in the Δ*parC* and Δ*parP* mutants was indistinguishable from that in the wild-type, as observed by confocal microscopy in cells expressing FliC^S267C^, a flagellin derivative that binds maleimide fluorescent dyes **(Pulido-Sánchez *et al*., 2025) (Fig. S2)**. We next analyzed the chemotactic response of the Δ*parC* and Δ*parP* mutants by performing chemical-in-plug chemotaxis assays **(Parales and Ditty, 2018)** using casamino acids as chemoattractant. Wild-type cells displayed motility halos biased towards the chemoattractant, showing an average chemotactic response index of 1.8 **(Fig. 1D and E)**. However, the Δ*parC* and Δ*parP* mutants exhibited symmetrical halos with response indexes close to 1, indicating that both ParC and ParP are required for the chemotactic behaviour of *P. putida*.

**Figure 1.**
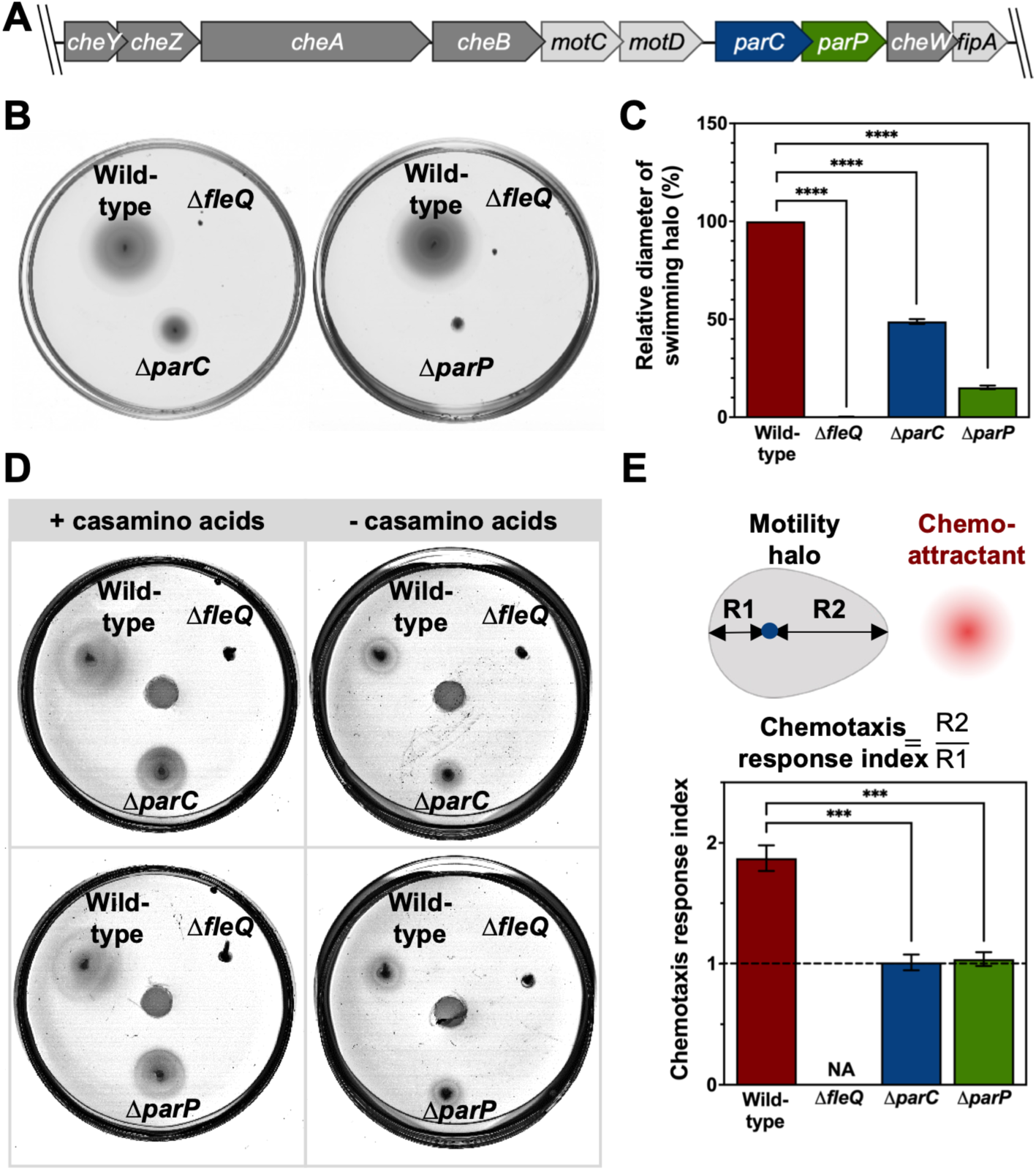
Swimming motility and chemotaxis assays of the Δ*parC* and Δ*parP* mutants. **A.** Cartoon showing the genomic context of the *parC* and *parP* genes in the *P. putida* KT2440 chromosome. Dark grey: chemotaxis machinery-encoding genes. Light grey: flagellar apparatus-related genes. **B.** Soft agar-based swimming motility assays of the Δ*parC* and Δ*parP* mutants. **C.** Motility halo diameters from the swimming motility assays. **D.** Soft agar-based chemical-in-plug chemotaxis assays towards an agarose plug with (left) or without (right) 0.2% casamino acids at the center of the plate. **E.** Cartoon depicting the calculation of the chemotaxis response index (top), and chemotaxis response indexes of the wild-type strain, and the Δ*parC* and Δ*parP* mutants when exposed to an agarose plug containing casamino acids (bottom). A dashed horizontal line indicates a chemotaxis response index of 1 (no response). (**B** and **D**) Each picture shows a representative swim plate out of at least three biological replicates. The wild-type strain and the Δ*fleQ* mutant were assayed in the same plates as positive and negative controls. (**C** and **E**) Columns and error bars represent the averages and standard deviations of at least three biological replicates. NA: not applicable. Stars designate *p*-values for Welch’s *t*-test in comparison with the wild-type strain (***:*p*<0.001; ****:*p*<0.0001).

### ParC and ParP form a complex with polar landmark FimV at the cell poles

Since ParC and ParP act as polar landmarks for the chemotaxis signaling arrays in *Vibrio* spp. **(Ringgaard *et al.,* 2018)**, we next assessed the intracellular localization of these proteins in *P. putida*. To this end, we constructed miniTn*7* derivatives to express ParC and ParP translational fusions to GFPmut3 under the salicylate-inducible P*sal* promoter. Insertion of the P*sal-parC-gfp* and P*sal-parP-gfp* transposon constructs in the chromosomes of the Δ*parC* and Δ*parP* mutants partially restored the swimming defects of the strains in the absence of salicylate, and salicylate addition improved complementation **(Fig. S3)**. Western blot assays using an anti-GFP antibody revealed a protein band compatible with the molecular weight of ParC-GFP (∼58 kDa), while a faint band migrating somewhat slower than expected (∼60 kDa) was obtained with ParP-GFP **(Fig. S4)**. ParP is a proline-rich (23/298) acidic (pI=4.76) protein featuring a long, unstructured region (**Ringgaard *et al*., 2018)**, all of which correlate with atypically slow migration on SDS PAGE **(Schram *et al*., 2019; Tiwari *et al*., 2019; Scheller *et al*., 2021).** In any case, positive complementation of the Δ*parP* strain as shown above suggests that, despite the observed anomaly, our ParP-GFP fusion retains ParP function.

ParC-GFP and ParP-GFP formed polar foci in 84% and 80% of the wild-type cells, respectively, with 60% and 12% of the cells exhibiting foci at both poles, respectively **(Fig. 2A and B)**. Fluorescent foci were never observed at non-polar locations. We analyzed the dynamics of ParC and ParP recruitment to the new cell pole during the cell cycle by examining the frequency of cells displaying bipolar distribution of the fluorescently labelled proteins as a function of cell length. Bipolar distribution of ParP-GFP was only observed in cells longer than ∼3.5 µm, and its frequency increased steadily with cell length **(Fig. 2C)**, mirroring our previous observations with the late flagellar protein FliC **(Pulido-Sánchez *et al*., 2025)**. These results suggest that ParP is recruited to the new cell pole concurrently with the final steps of flagellar assembly before cell division. In contrast, bipolarly distributed ParC-GFP accumulated early in the cell cycle to peak in cells of intermediate size and decline again in longer, predivisional cells **(Fig. 2C)**. Notably, ∼50% of the shorter, postdivisional cells displayed no ParC foci, although this frequency decreased with growth to a minimum <10% in cells longer than ∼3.5 µm **(Fig. 2D)**. These observations suggest that in cells bearing ParC at the old (flagellated) pole, the new pole is populated with ParC units at the expense of the old pole. While this leads to a transient bipolar distribution, eventually the old pole becomes vacant, resulting in ParC association only at the new pole. This inevitably leads to asymmetric segregation, with only one of the daughter cells inheriting ParC nucleated at a single pole.

**Figure 2.**
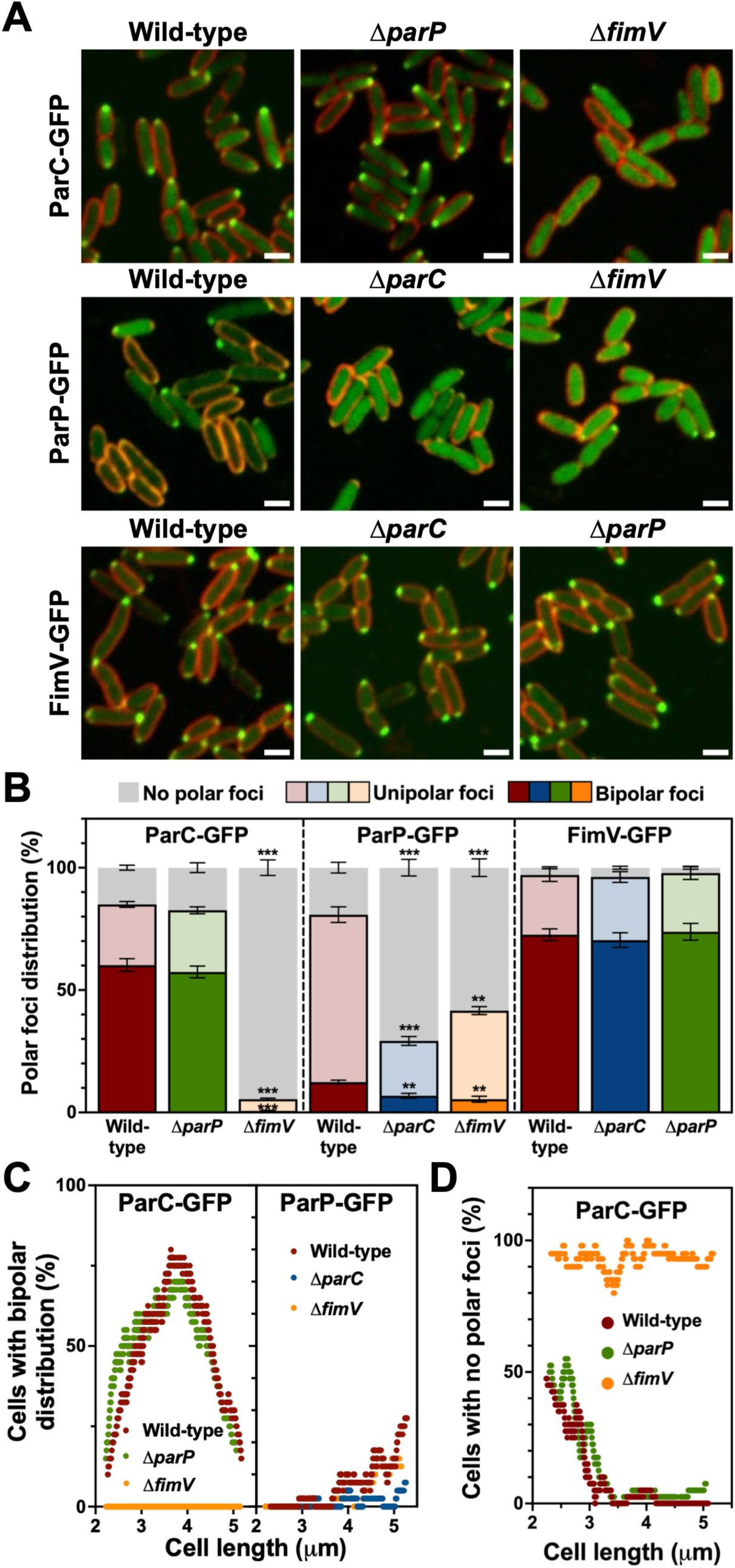
Intracellular location of ParC, ParP and FimV. **A.** Confocal microscopy images of wild-type, Δ*parC,* Δ*parP* and Δ*fimV*.cells expressing P*sal-parC-gfp*, P*sal-parP-gfp* or P*sal-fimV-gfp* (green) in the absence (FimV) or the presence (ParC and ParP) of 2 mM salicylate. FM^TM^ 4-64 was used as membrane stain (red). Images are shown as the maxima projections of seven Z-sections of the green channel, merged with the red channel showing the cell contour at the focal plane. Scale bar: 2 µm. **B.** Frequency of cells bearing bipolar, unipolar o no polar ParC-GFP, ParP-GFP or FimV-GFP foci. Columns and error bars represent the averages and standard deviations of at least three separate replicates. Stars designate *p*-values for Welch’s *t*-test (**:*p*<0.01; ***:*p*<0.001) in comparisons with the wild-type strain. **C.** Frequency of cells displaying bipolar distribution of ParC-GFP or ParP-GFP foci *vs.* cell length. **D.** Frequency of cells displaying no polar ParC-GFP foci *vs.* cell length. (**C** and **D**) Plots represent frequencies in overlapping sets of 40 consecutive segmented, length-sorted cells plotted against the average length of each set (n = 500 cells).

Next, we examined the impact of ParC on ParP location and vice versa. While deletion of *parP* did not substantially alter the polar distribution of ParC-GFP **(Fig. 2A and B)** or its dynamics in the cell cycle **(Fig. 2C and D)**, Δ*parC* cells displayed a diffuse distribution of ParP-GFP in the cytoplasm, accompanied by localized foci at one or both cell poles in only 29% of the population **(Figs. 2A and B).** Again, ParC-GFP and ParP-GFP foci were never detected at non-polar locations. Bipolar ParP-GFP distribution was restricted to a small percentage of very long, predivisional cells in this background **(Fig. 2C)**. These results indicate that ParC is required for efficient recruitment of ParP to the new cell pole.

In *V. cholerae* and *V. parahaemolyticus*, HubP determines the polar localization of ParC **(Yamaichi *et al*., 2012)**. Lack of the HubP ortholog in *P. putida*, FimV **(Pulido-Sánchez *et al*., 2025)** provoked a defect in chemotaxis, as shown by a significant reduction in the chemotactic response index towards the assayed chemoattractant **(Fig. S5)**. Accordingly, we next tested the possible involvement of FimV in the polar localization of ParC and ParP. To this end, we inserted the P*sal-parC-gfp* and P*sal-parP-gfp* transposons in the chromosome of the Δ*fimV* mutant, as well as a previously constructed P*sal-fimV-gfp* transposon **(Pulido-Sánchez *et al*., 2025)** into the Δ*parC* and Δ*parP* mutants, and assessed the intracellular distribution of the fluorescent proteins **(Fig. 2)**. While the intracellular distribution of FimV-GFP was not altered in the absence of ParC or ParP **(Fig. 2A and B)**, deletion of *fimV* caused a major disruption of the polar localization of ParC-GFP and ParP-GFP. In the absence of FimV the frequency of cells showing polar ParC-GFP foci dropped to ∼5%, with a residual population (<1%) showing foci at both cell poles **(Fig. 2A and B)**. While FimV was not essential for polar localization of ParP-GFP, the frequency of cells bearing unipolar or bipolar polar foci was decreased by half in the Δ*fimV* mutant **(Fig. 2A and B)**. In addition to polar foci, we note that Δ*fimV* cells displayed diffuse ParP-GFP fluorescence in the cytoplasm **(Fig. 2A)**, reminiscent of the ParP-GFP distribution in the Δ*parC* strain **(Fig. 2A)**. Because of the shared location of the flagellar and chemotaxis machineries and their related functions, we explored the possibility of cross-talk between ParC and ParP and the regulators of flagellar location and number FlhF and FleN. The intracellular distribution of the fluorescently labeled FlhF-GFP and GFP-FleN **(Pulido-Sánchez *et al*., 2025)** was not altered by the absence of ParC or ParP (**Fig. S6**). Similarly, deletion of *flhF* or *fleN* did not alter the intracellular distribution of ParC-GFP or ParP-GFP (**Fig. S7**), suggesting that both recruitment pathways are independent from each other.

### FimV, ParC and ParP interact directly with each other

We next assessed the *in vivo* interactions between the three proteins by way of bacterial adenylate cyclase-based two-hybrid (BACTH). To this end, the T18 and T25 adenylate cyclase domains were fused to the C-terminal ends of FimV, ParC and ParP in plasmids pUT18 and pKT25. Combinations of T18 and T25 fusions were transformed into the reporter *E. coli* strain BTH101, and interaction was assessed visually as growth and blue colonies on minimal lactose medium containing X-gal, and quantified by means of β-galactosidase assays **(Fig. 3)**. We observed high β-galactosidase levels in the FimV-FimV and FimV-ParC pairs, and modest but significant β-galactosidase levels with the FimV-ParP and ParC-ParP pairs, suggesting that interaction with the polar landmark protein FimV may contribute to polar recruitment of ParC, while interaction with FimV and ParC may contribute to polar recruitment of ParP. The ParC-ParC and ParP-ParP pairs yielded β-galactosidase activity levels indistinguishable from the negative control, indicating that our assay failed to detect dimerization of these two proteins, as previously described **(Ringgaard *et al*., 2014)**.

**Figure 3.**
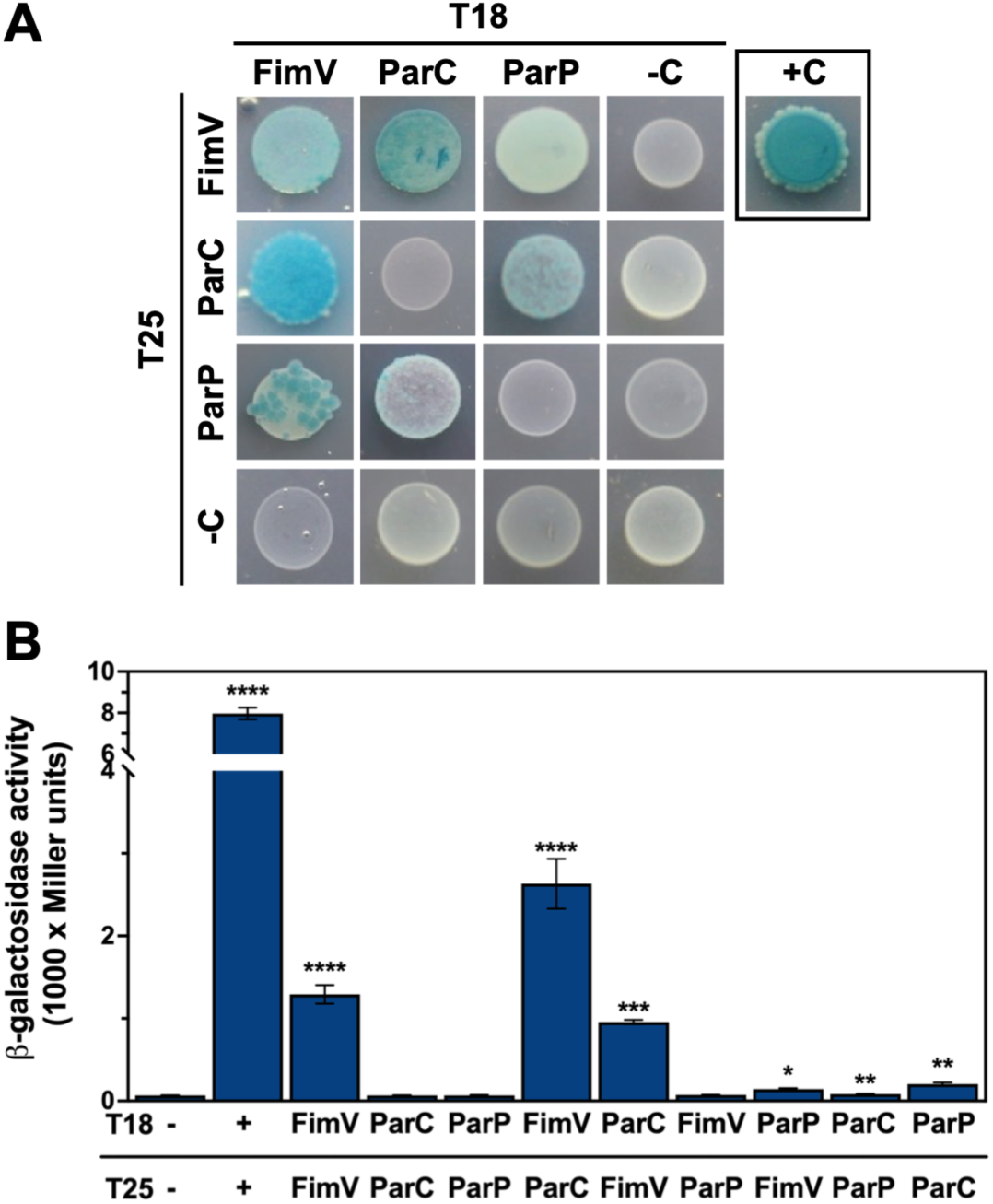
Bacterial two-hybrid assays between FimV, ParC and ParP. **A.** Interaction drop-assays between FimV, ParC and ParP proteins fused to adenylate cyclase fragments T18 and T25 in the the *E. coli* BTH101 reporter strain. Colonies were grown in minimal medium supplemented with lactose and X-gal. Colony growth and blue colour denote positive interaction between each pair of hybrid proteins. Co-transformants expressing pUT18 and pKNT25 empty vectors, or pUT18-zip and pKNT25-zip were used as negative and positive controls, respectively. Each picture shows a representative colony out of at least nine biological replicates. **B.** Quantification of the interaction strength by β-galactosidase assays. Columns and error bars represent the averages and standard deviations of at least three biological replicates. Stars designate *p*-values for Welch’s *t*-test in comparisons with the negative control (NS: not-significant; *:*p*<0.05; **:*p*<0.01; ***:*p*<0.001; ****:*p*<0.0001).

We explored further the interactions between FimV, ParC and ParP by AlphaFold 3-driven modeling of protein complexes **(Abramson *et al*., 2024) (Fig. S8 and Supplementary Data 1)**. FimV is a transmembrane protein with its N-terminus oriented to the periplasm and its C-terminus located in the cytoplasm **(Schmidt *et al*., 2026)**. Comparative analysis of five independent models of the interaction between the cytoplasmic FimV^410-911^ region with ParC robustly defined contacts between FimV residues 589-599 and three ParC regions (residues 103-114, 134-143 and 157-175), located at and around the Walker B motif of the ATP binding and hydrolysis domain **(Ringaard *et al*., 2011)**. In addition, FimV residues 855-890 interacted with ParC 204-217 and 240-254 **(Fig. S8A and B)**. Similarly, models of FimV^410-911^ and ParP interaction displayed a consistent contact surface between FimV residues 714-736 and ParP 211-216 and 253-274 residues located at the CheW-like domain **(Fig. S8C and D)**. Finally, models of ParC interaction with ParP consistently predicted interactions of ParC residues 103-110, 133-143 and 164-175 and ParP 15-25 **(Fig. S8E and F)**. Taken together, our *in vivo* experimental and *in silico* modelling results strongly suggest that the FimV>ParC>ParP polar recruitment cascade is supported by direct protein-protein interactions.

### Polar targeting of CheA is mediated by FimV, ParC and ParP

To address how FimV, ParC and ParP contribute to the recruitment of the chemotaxis proteins, we used the histidine kinase CheA as reporter. To assess the intracellular localization of CheA, we engineered a miniTn*7* derivative to produce a translational fusion of this protein to GFPmut3 from its natural P*cheA* promoter. Western blot assays using an anti-GFP antibody revealed a protein band compatible with the molecular weight of CheA-GFP (∼108 kDa) **(Fig. S4)**. Complementarily, we constructed a KT2442 derivative bearing a complete in-frame deletion of the *cheA* gene to assess functional complementation of the fusion protein. The Δ*cheA* mutant did not spread from the inoculation point in soft agar-based swimming in the standard duration of the experiment (12 h) (**Fig. S9A**), but displayed a small motility halo after 48 h incubation **(Fig. S9B)** and single-cell motility tracking revealed that the mutant cells are fully motile, but largely unable to change swimming direction as required for the chemotactic response **(Fig. S9C)**. Expression of P*cheA-cheA-gfp* restored the ability to display swimming motility halos in the Δ*cheA* mutant (**Fig. S9A**), indicating that the fusion protein retains CheA function.

By using confocal microscopy, we observed that wild-type cells bearing the P*cheA-cheA-gfp* transposon displayed heterogeneous diffuse fluorescence in the cytoplasm, occasionally accompanied by polar foci (**Fig. S10**). In contrast, polar CheA-GFP foci were evident in most cells of the Δ*cheA* mutant, with little residual fluorescence in the cytoplasm **(Fig. 4A)**. We attribute these differences to an apparent competitive advantage of the native CheA protein in its association to the cell poles. Consequently, we further performed CheA-GFP microscopy imaging in Δ*cheA* strains. The vast majority (∼99%) of Δ*cheA* cells exhibited polar CheA-GFP foci **(Fig. 4A and B)** and ∼20% displayed a bipolar distribution. Deletion of *parC*, *parP* or *fimV* in this background only caused a small decrease (5-12%) in the frequency of polar foci, and only the Δ*fimV* mutant displayed somewhat higher (∼30%) frequency of the bipolar distribution, indicating that none of these proteins are strictly required for the recruitment of CheA to the cell poles **(Fig. 4A and B)**. However, a general decrease in the fluorescence intensity of the polar CheA-GFP foci was also noted in the absence of FimV, ParC or ParP **(Fig. 4C)**. This was accompanied by 60 and 34% increased frequency of cells displaying ectopic CheA-GFP foci at non-polar locations in the Δ*parC* Δ*fimV* mutants, respectively **(Fig. 4D)**, suggesting that these two proteins are required for specific nucleation at the cell poles. In contrast, ectopic foci were not more prevalent in the Δ*parP* mutant **(Fig. 4D)**. Instead, a higher intensity of diffuse fluorescence in the cytoplasm was found in this background **(Fig. 4A)**, suggesting while ParP may be required to stabilize CheA within the cluster lattice and prevent its release to the cytoplasm. When examining the dynamics of bipolar CheA-GFP distribution relative to cell length in the Δ*cheA* mutants, we observed that the frequency increased steadily with cell length to a maximum ∼80% in long, predivisional cells **(Fig. 4E)**, suggesting that new CheA molecules are recruited to the new cell pole while others remain permanently associated to the old cell pole, as previously observed with flagellar proteins **(Pulido-Sánchez *et al*., 2025)**. This pattern is unaltered in the absence of FimV. However, deletion of *parC* or *parP* resulted in CheA-GFP accumulation at the new pole stalling at 40-50% bipolar distribution in cells longer than ∼4 µm **(Fig. 4E).** In addition, CheA accumulation was more prevalent in shorter cells in the absence of ParC. These results suggest that CheA can spontaneously locate at the cell poles, but this association is temporary and in a dynamic equilibrium with the cytoplasmic CheA pool. At the early stages of the cycle ParC limits CheA interaction with the cell pole, but polar recruitment of ParP at the later stages boosts permanent association of CheA with the cell poles, likely by promoting its assembly into the chemotaxis arrays. We also assessed the impact of CheA on the location of ParC-GFP and ParP-GFP **(Fig. S11)**, but no changes in the intracellular distribution of both proteins were evident, suggesting that CheA does not contribute to the polar location of ParC or ParP.

**Figure 4.**
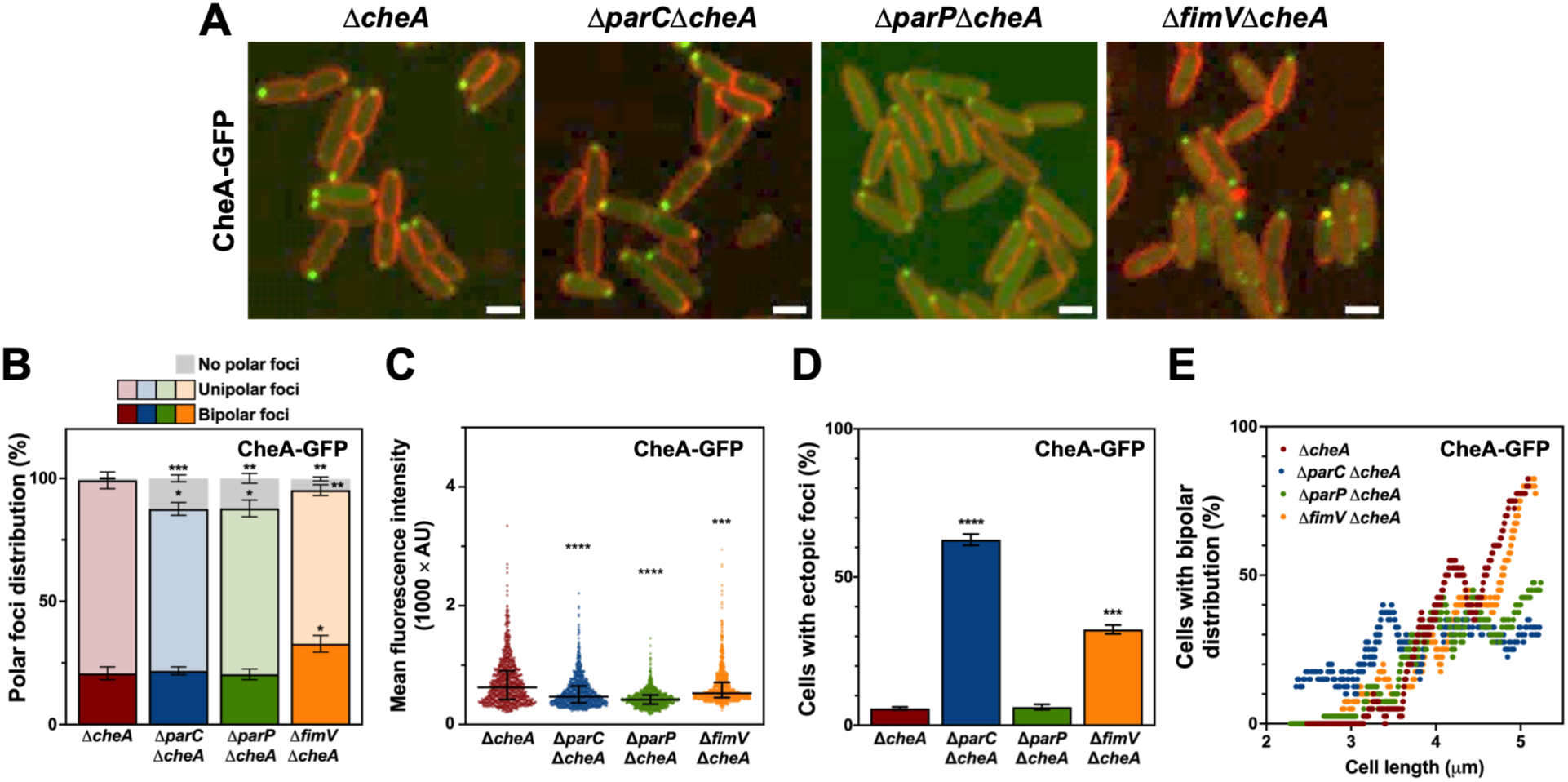
Intracellular location of CheA. **A.** Confocal microscopy images of Δ*cheA*, Δ*parC*Δ*cheA*, Δ*parP*Δ*cheA* and Δ*fimV*Δ*cheA* cells expressing P*cheA-cheA-gfp* (green). FM^TM^ 4-64 was used as membrane stain (red). Images are shown as the maxima projections of seven Z-sections of the green channel, merged with the red channel showing the cell contour at the focal plane. Scale bar: 2 µm. **B.** Frequency of cells bearing bipolar, unipolar o no polar CheA-GFP foci. Columns and error bars represent the averages and standard deviations of at least three separate replicates. **C.** Mean fluorescence intensity of polar CheA-GFP foci. Points correspond to individual fluorescence intensity values, bars denote the median and the interquartile interval (n = 500 cells). **D.** Frequency of cells bearing non polar CheA-GFP foci. Columns and error bars represent the averages and standard deviations of at least three separate replicates. **E.** Frequency of cells displaying bipolar distribution of CheA-GFP foci *vs.* cell length. The plot represents frequency of bipolar distribution in overlapping sets of 40 length-sorted cells plotted against the average length of each set (n = 500 cells). Stars designate *p*-values for Welch’s *t*-test (**B, D**), or the pairwise Mann-Whitney U test (**C**) in comparisons with the Δ*cheA* strain (ns: not significant; *:*p*<0.05; **=*p*<0.01;***:*p*<0.001; ****:*p*<0.0001).

### ParC, ParP and CheA are required for polar clustering of the chemoreceptor Aer2

To evaluate the role of the ParC/ParP system in the localization of the chemotaxis chemoreceptors in *P. putida*, we engineered a translational fusion to *yfp* at the 3’ end of the *aer2* chemoreceptor-encoding gene at its genomic *locus* in the wild-type strain and the Δ*parC*, Δ*parP* and Δ*fimV* mutants. *P. putida* cells bearing this fusion protein were previously shown to display polar fluorescent foci, consistent with the location of Aer2 as part of the chemosensory arrays **(Sarand *et al*., 2008)**. Confocal microscopy revealed that in wild-type cells, Aer2-YFP was diffusely distributed at the cell contour (**Figs. 5A and B and S12**), but accumulated as foci at a single pole in 90-95% of the cells, with 1-2% displaying foci at both poles **(Fig. 5C)**. Bipolar distribution of Aer2-YFP was only observed in cells longer than ∼3.5 µm, and its frequency increased steadily with cell length **(Fig. S13)**, as shown above for ParP **(Fig. 2C)**. In contrast, polar accumulation of Aer2-YFP was not observed in the Δ*parC* and Δ*parP* strains **(Fig. 5A, B and C)**, indicating that the formation of polar chemosensory arrays is strictly dependent on the ParC/ParP system. We did not observe major changes in the cellular distribution of Aer2-YFP in the Δ*fimV* mutant, and only a small increase in the frequency of bipolar Aer2-YFP foci was evident in this background **(Figs. 5A**, **B and C)**. Deletion of FimV however caused a decrease in the intensity of the Aer2-YFP foci, suggesting that it may directly or indirectly stimulate the recruitment or stabilize the association of the chemoreceptors to the cell poles.

**Figure 5.**
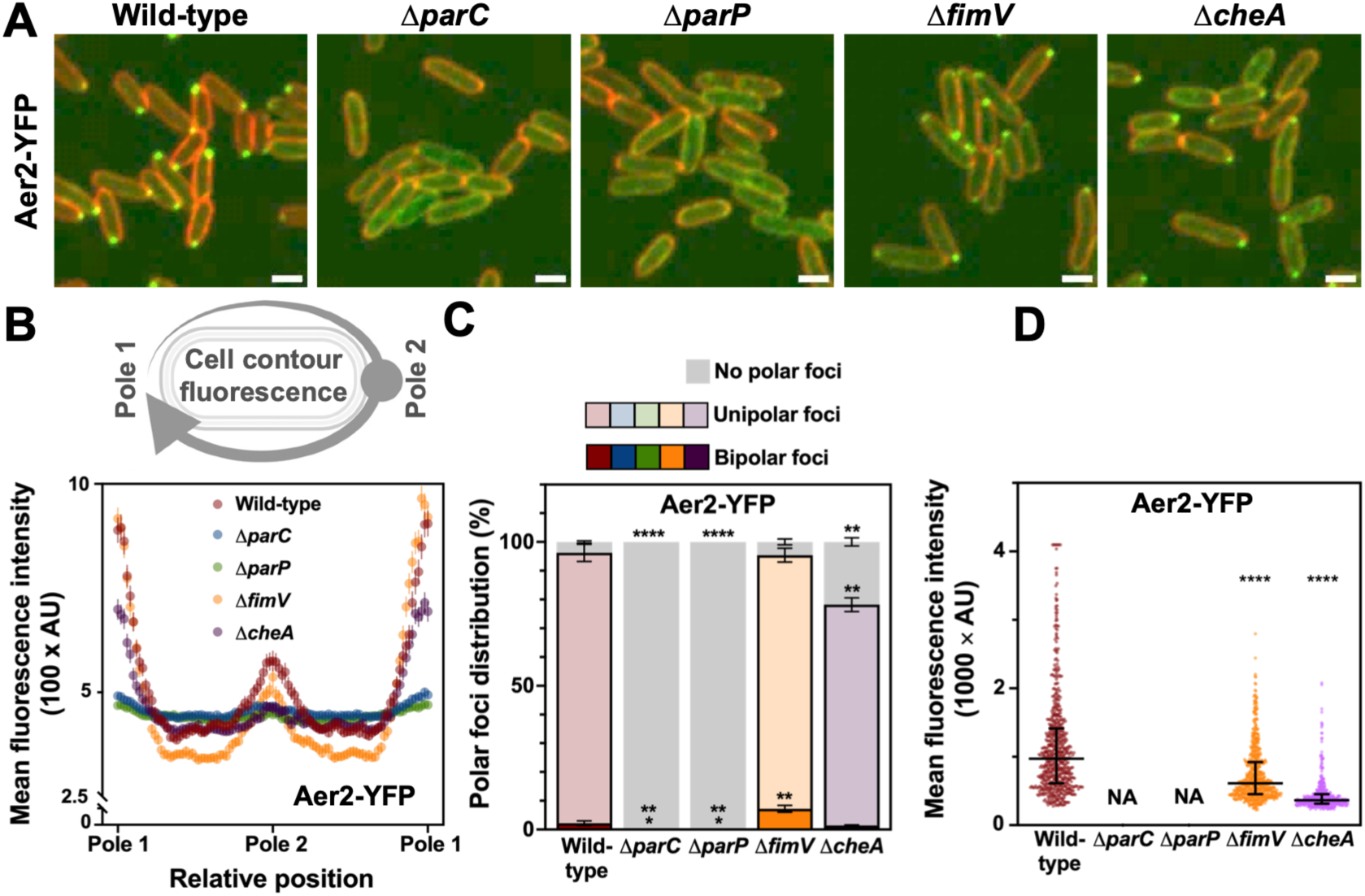
Intracellular location of MCPs. **A.** Confocal microscopy images of wild-type, Δ*parC*, Δ*parP*, Δ*fimV* and Δ*cheA* cells expressing *aer2-yfp in locus* (green). FM^TM^ 4-64 was used as membrane stain (red). Images are shown as the maxima projections of seven Z-sections of the yellow channel, merged with the red channel showing the cell contour at the focal plane. Scale bar: 2 µm. **B.** Distribution of and Aer2-YFP fluorescence along the cell contour. The x-axis displays the relative position along the cell contour following the path depicted by the upper scheme: the axis origin represents the cell pole with the highest fluorescence intensity (pole 1), mid-axis represents the pole with the lowest fluorescence intensity (pole 2), and right-end represents again pole. Points and error bars represent the averages and standard deviations of at least three separate replicates (n = 500 cells). **C.** Frequency of cells bearing bipolar, unipolar o no polar Aer2-YFP foci. Columns and error bars represent the averages and standard deviations of at least three separate replicates (n = 500 cells). **D.** Fluorescence intensity of polar Aer2-YFP foci. Points correspond to individual fluorescence intensity values, bars denote the median and the interquartile interval. Stars designate *p*-values for Welch’s *t*-test (**C**), or the pairwise Mann-Whitney U test (**D**) in comparisons with the wild-type strain (*:*p*<0.05; **=*p*<0.01;***:*p*<0.001; ****:*p*<0.0001).

Similarly, we examined whether CheA plays a role in the polar location of the chemoreceptors. Aer2-YFP was still able to locate at the cell contour and form polar foci in the absence of *cheA* **(Fig. 5A and B)**. Fluorescence at the cell contour was similar to the wild-type strain **(Fig. 5B)**, and the frequency of cells bearing polar foci was somewhat diminished **(Fig. 5C)**. In addition, we observed a substantial reduction in the fluorescence intensity of the polar foci in the mutant background **(Fig. 5B and D),** suggesting that CheA is not strictly required for polar location, but stimulates polar clustering of the chemoreceptors at the chemosensory arrays.

As above, we performed AlphaFold 3-driven modelling of ParP interaction with CheA and Aer2 **(Fig. S14 and Supplementary Data 1)**. Models consistently predicted contacts between residues 153-174 and 225-244 at the ParP CheW-like domain and 367-394 at the Aer2 protein contact tip **(Fig. S14A and B)**. Similarly, ParP was predicted to interact with CheA by way of residues 253-274. The ParP interaction surface in CheA was not consistent across structural models. However, contacts in CheA recurrently mapped to at least one of three 19-bp repeat regions spanning positions 196 and 295, with the central repeat being the most common occurrence. Repeats 1 and 2 displayed the identical sequence DEITDAEFESLLDQLHGKG, while repeat 3 was degenerate, but preserved the core EFExLLD motif **(Fig. S14C and D)**. The interaction regions predicted are generally consistent with those experimentally determined in *Vibrio* spp., suggesting that they may also represent genuine sites of protein-protein interaction in *P. putida*.

## DISCUSSION

Despite their lack of internal compartments, prokaryotic cells are highly organized, with related functional elements often recruited to specific cellular locations to achieve maximum efficiency **(Surovtsev and Jacobs-Wagner, 2018; Rudner and Losick, 2010)**. The delivery of the components of the bacterial chemosensory systems to particular regions in the cell envelope and their assembly into intricate arrays containing multiple sensory elements along with the signal transduction system required to translate the signals detected into a coherent behavior is but one example of such elaborate intracellular organization **(reviewed by Jones and Armitage, 2015)**. Here we present evidence of the involvement of the polar landmark proteins FimV, ParC and ParP in the stable recruitment of the components of the chemosensory apparatus to the cell poles of *P. putida* to form functional chemosensory arrays in the vicinity of the polar flagellar apparatus during its assembly. By coordinately assembling the flagella and associated chemosensory machinery in the timespan of one cell cycle, polarly flagellated bacteria ensure the inheritance of functional motility and chemotaxis systems by both daughter cells after cell division.

Our results demonstrate that the polar clustering required for full functionality of the chemosensory arrays is driven by the action of three polar landmark proteins: FimV, ParC and ParP. We have shown that all three proteins are required for wild-type chemotaxis **(Fig. 1 and S5)**, and FimV is responsible for recruitment of ParC to the cell poles, while both FimV and ParC contribute to polar location of ParP **(Fig. 2)**. The temporal dynamics of the polar location of the three proteins lends additional support to this recruitment hierarchy. Firstly, FimV is recruited to the midcell region around the time of cell division to remain associated to the new cell pole of the newborn cells **(Pulido-Sánchez *et al*., 2025)**. Secondly, ParC is recruited to the cell pole after cell division and accumulates during the first half of the cell cycle in a FimV-dependent fashion **(Fig. 2C)**. Finally, ParC-dependent polar accumulation of ParP only occurs at the late stages of the cycle, **(Fig. 2C)**.

On the surface, the FimV>ParC>ParP cascade appears to replicate the HubP>ParC>ParP hierarchy thoroughly characterized in *V. cholerae* and *V. parahaemolyticus* **(Ringgaard *et al*., 2011; Yamaichi *et al*., 2012; Ringgaard *et al*., 2014; Alvarado *et al*., 2017)**. However, we note several discrepancies between both models. Firstly, the impact of HubP on ParC polar accumulation is relatively minor in *Vibrio* spp., as most Δ*hubP* cells still display polar ParC foci, and the most distinct feature documented is the presence of additional foci at non-polar locations **(Yamaichi *et al*., 2012)**. In contrast, ParC polar location is largely dependent on the presence of FimV in *P. putida* **(Fig. 2A-D)** and mislocalized ParC does not produce ectopic foci, but is diffusely distributed in the cytoplasm. A second difference is observed in the dynamics of ParC accumulation. In *V. cholerae*, ParC is recruited and becomes permanently bound to the new cell pole shortly after cell division **(Ringgaard *et al*., 2011)**. In contrast, the frequency of ParC bipolar distribution in *P. putida* peaks in cells of intermediate length to drop again in longer, predivisional cells **(Fig. 2C)**. ParC release results in a vacant pole that is transmitted to one of the daughter cells **(Fig. 2D)**. This result indicates that stable ParC association with the cell pole beyond the assembly stage is not essential for normal function of the chemotaxis system, at least in *P. putida*. Finally, ParC and ParP are simultaneously recruited to the new cell pole in *Vibrio* spp. **(Ringgaard *et al*., 2014)**, while a distinct gap between the timing of ParC and ParP recruitment is evident in *P. putida*, to the extent that ParP accumulation occurs concurrently to ParC disengagement **(Fig. 2C)**. This delay is not due to differentially regulated synthesis of both proteins, as they are both produced ectopically in this work, and suggests that a physiological regulatory switch operating at the posttranslational level may preclude immediate incorporation of ParP to the new ParC-populated cell pole.

Some of the contrasting observations noted above may originate from the different mechanisms of ParC recruitment in *Vibrio* spp. and *P. putida*. In *V. cholerae*, recruitment of ParC to the new pole is the result of the redistribution of molecules bound to the old pole by way of a diffusion-and-capture mechanism to achieve a bipolar distribution at the equilibrium in the longer, predivisional cells **(Ringgaard *et al*., 2011)**. We propose that a diffusion-and-capture mechanism is also responsible for ParC recruitment in *P. putida*. However, recapture at the old pole is somehow prevented, and ParC progressively accumulates at the new pole as it becomes depleted from the old pole, resulting in a transient bipolar distribution that evolves into a unipolar distribution with ParC located only at the new pole at the end of the cycle. Multiple lines of evidence suggest that ParC does not interact directly with HubP in *V. cholerae*, and HubP-dependent recruitment is dependent on an as of yet uncharacterized linker protein. In addition, ParC recruits ParP by way of protein-protein interactions and ParP promotes ParC retention at the cell pole **(Ringgaard *et al*., 2011)**. The fact that a substantial amount of ParC is present at the cell poles in the Δ*hubP* mutant suggests that the combined action of the linker protein and ParP may be sufficient to support polar location of ParC in the absence of HubP, although preference for the cell poles is increased further in its presence. In contrast, in *P. putida* FimV, ParC and ParP interact directly with each other *in vivo* in BACTH assays **(Fig. 3)**, and AlphaFold modeling predicts interactions with FimV and ParP to occur at the same interface in ParC: three distinct regions located in the vicinity of the Walker B motif **(Fig. S1, S8A and B and S14A and B)**. Models also predict interaction of these regions with a discrete region in the proximal section of the cytoplasmic C-terminal domain of FimV. An additional interaction surface is predicted between the C-terminal domain of ParC and the conserved FimV motif at the distal C-terminal end of FimV **(Fig. S8)**. However, we recently showed that removal of the FimV motif does not affect the chemotactic behavior of *P. putida* **(Schmidt *et al*., 2026)**, suggesting that the latter interaction is likely not relevant to ParC recruitment. We propose that, rather than stabilizing ParC at the cell pole, ParP competes with FimV for their shared interaction region in ParC. As a result, ParC tether to FimV is weakened or abolished in the presence of ParP and its interaction with the cell pole is compromised, leading to its eventual dissociation. The validity of the modeling results obtained and their relevance to this hypothesis will be challenged experimentally in the near future.

We have also examined the contribution of FimV, ParC and ParP to the polar location of the chemotaxis machinery components. The kinase/phosphotransferase protein CheA is part of the signal transduction pathway of chemotaxis and a cytoplasmic component of the chemosensory arrays **(Muok *et al*., 2020; Wadhams and Armitage, 2004)**. Aer2 is a chemoreceptor similar to *E. coli* Aer **(Sarand *et al*., 2008)**. Our results show that CheA and Aer2 accumulate at polar locations in the vast majority of wild-type cells **(Fig. 4A and B and 5A and C)**. Similarly to ParP, CheA and Aer2 shifted their distribution from unipolar to bipolar with increasing cell length **(Figs. 4E and S10)**, suggesting permanent association to the old (flagellated) pole and progressive occupancy of the new (non-flagellated) pole in the timespan of one cell cycle **(Pulido-Sánchez *et al*., 2025)**. While the overall ability of CheA to associate to the cell poles was not greatly affected by the presence of FimV, ParC or ParP **(Fig. 4A and B)**, the full extent of CheA accumulation was not attained in their absence **(Fig. 4D)**. Polar localization of CheA was inhibited by ParC in shorter cells, but was stabilized by ParC-dependent recruitment of ParP at later stages **(Fig. 4E)**. We propose that CheA can spontaneously associate to the cell poles, but the interplay between ParC and ParP promotes timely, permanent association of CheA with the new pole specifically in long, predivisional cells, as shown in *Vibrio* spp. **(Ringgaard *et al*., 2014; Alvarado *et al*., 2017)**. In contrast, polar nucleation of Aer2 was strictly dependent on the presence of ParC and ParP **(Fig. 5A-C)**, and the extent of its polar accumulation was greatly increased by CheA **(Fig. 5D)**. The temporal and spatial coincidence of ParP, CheA and the MCPs at the cell poles and their dependence relationships for recruitment and stabilization are fully consistent with the recruitment and assembly model proposed for *V. cholerae* and *V. parahaemolyticus*. In these organisms, ParC recruits ParP by direct interaction with its N-terminal ParC-binding domain. Then, ParP interacts with the Localization and Inheritance Domain (LID) of CheA and the protein interaction tips of the MCPs by way of its C-terminal CheW-like Array Integration and Formation (AIF) domain to become an integral part of the chemosensory arrays **(Ringgaard *et al*., 2014; Alvarado *et al*., 2017)**. As a result, ParP stabilizes the polar association of CheA, prevents diffusion of the chemoreceptors in the cell membrane and promotes their assembly into the polar chemosensory arrays **(Ringgaard *et al*., 2014; Alvarado *et al*., 2017; Ringgaard *et al*., 2018)**. AlphaFold 3-based modeling of these interactions between the *P. putida* proteins accurately predicts ParP contact surfaces with ParC, CheA and Aer2 **(Figs. S1 and S14)** that align with the experimentally determined interaction sites in *Vibrio* spp. **(Ringgaard *et al*., 2014; Alvarado *et al*., 2017)**. Similarly, the predicted ParP interaction region at the cytoplasmic tip of Aer2 overlapped the conserved QTNLLALNAAI motif described in *Vibrio* spp. **(Fig. S14A and B)**. Finally, CheA contacts with ParP were less conserved, but often mapped to at least one of three copies of a 19-bp degenerate repeated sequence featuring the conserved core motif EFExLLD **(Fig. S14C and D)**. These repeats are exclusive to CheA proteins from organisms containing ParC and ParP **(Fig. S15)** and overlap the LID domain shown to be sufficient for *in vivo* interaction with ParP in *V. parahaemolyticus* **(Ringgaard *et al*., 2014)**. Notwithstanding the lack of experimental evidence for the above interactions, the high degree of coincidence with the experimental results obtained in *Vibrio* spp. suggests that the AlphaFold-based models likely have a high predicted value worth testing experimentally.

Taken together, the evidence presented supports the notion that the ParC/ParP system is accountable for the recruitment and assembly of the chemosensory arrays at the cell poles in *P. putida*, as previously reported in *Vibrio* spp. **(Ringgaard *et al*., 2014; Alvarado *et al*., 2017; Ringgaard *et al*., 2018)**, although some of the mechanisms involved represent a departure from the *Vibrio* systems, as discussed above. We propose that FimV captures cytoplasmic ParC at the new cell pole where flagella assembly is taking place during the early stages of the cell cycle **(Fig. 6)**, while CheA associates and dissociates spontaneously from the cell pole. This association dynamics may be supported by transient interactions with MCPs diffusing at the cell membrane and is limited by the presence of ParC. At a later stage, ParC facilitates polar targeting of ParP concomitantly and in turn, ParP directs polar assembly of the chemosensory arrays by integrating in the lattice and stabilizing the chemosensory components. Our model highlights the driving role of the multiple protein-protein interactions displayed by ParP in the assembly of the chemosensory arrays, as described in *V. parahaemolyticus* and *V. cholerae* **(Ringgaard *et al*., 2014; Alvarado *et al*., 2017)**. In summary, our results strongly support a model in which the assembly of the chemotaxis arrays arises from the collaborative interactions between FimV, ParC and ParP and the chemoreceptors and chemotaxis signal transduction proteins.

**Figure 6.**
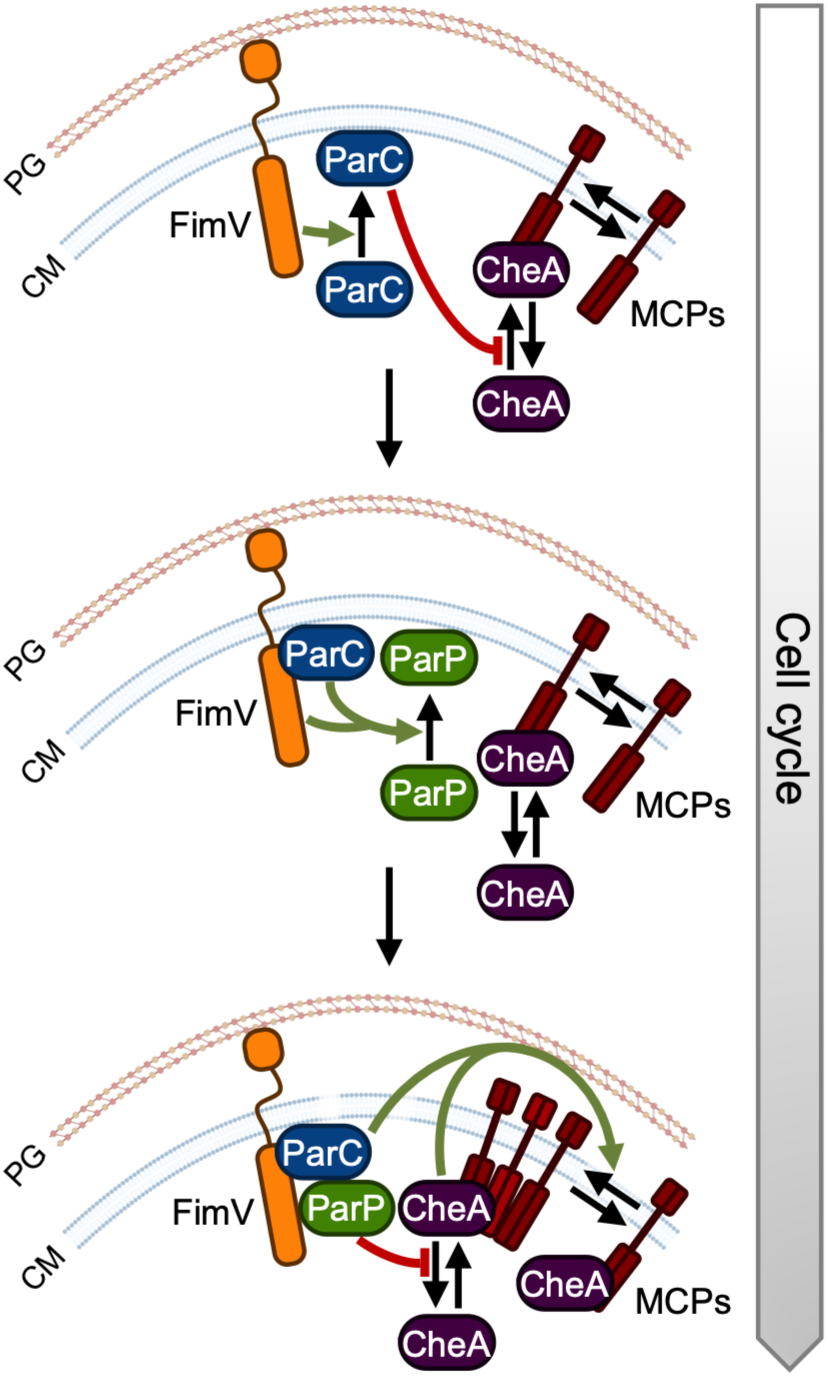
Working model of the polar localization of the chemotaxis arrays in *P. putida*. Cartoon illustrating the hierarchical interactions between FimV, ParC and ParP that mediate localization of the MCPs and CheA at the new cell pole during the cell cycle. Green arrows and T-shaped end red lines denote positive and negative effects, respectively. CM: cell membrane. PG: peptidoglycan.

Positioning of the chemosensory arrays at or near the cell poles has been vindicated as a safe means to guarantee inheritance of chemotaxis functionality by both offspring upon cell division **(Jones and Armitage, 2015)**. The elegant mechanisms previously described for *Vibrio* spp. **(Ringgaard *et al*., 2011; Yamaichi *et al*., 2012; Ringgaard *et al*., 2014; Alvarado *et al*., 2017)** and *S. putrefaciens* **(Rossmann *et al*., 2015),** and herein for *P. putida* attain this goal by coupling the *de novo* assembly of chemotaxis arrays at the new cell pole to the progress of the cell cycle. Because both *Vibrio* and *Pseudomonas* spp. bear polar flagella, the chemosensory arrays are located in close vicinity to the flagellar apparatus. The findings reported here mirror our previous work with the *P. putida* polar flagella **(Pulido-Sánchez *et al*., 2025)**. Flagella are also sequentially assembled at the new pole in the duration of a cell cycle in preparation for cell division. Both processes are initiated by FimV and involve the recruitment of a MinD/ParA family ATPase (i.e., ParC and FleN), followed by an additional protein (ParP and FlhF), which orchestrates the polar assembly of the structural components of each complex. The correct and timely operation of these mechanisms secures the inheritance of a functional chemotaxis-driven flagellar apparatus by both daughter cells after cell division.

## ACKNOWLEDGMENTS

We wish to thank Victoria Shingler for providing the pVI812 plasmid, Katherina García from the Microscopy Facility at CABD (Sevilla, Spain) for excellent technical assistance, and Kai Thormann for thoughtful advice and discussion. This work was supported by grants PGC2018–097151-B-I00, PID2021–126121-NB-I00 and CEX2020–00108-M from the Spanish Ministerio de Ciencia, Innovación y Universidades/Agencia Española de Investigación (MCIN/AEI) and the European Regional Development Fund “A way of making Europe” (ERDF), and by a predoctoral Formación de Profesorado Universitario contract 19/02899 of the Spanish Ministerio de Educación y Formación Profesional, awarded to M P-S.

## AUTHOR CONTRIBUTIONS

Marta Pulido-Sánchez: Data curation, formal analysis, investigation, methodology, writing - review and editing

María Zapata-Cruz: Formal analysis, investigation, methodology Antonio Leal-Morales: Formal analysis, investigation, methodology

Aroa López-Sánchez: Conceptualization, methodology, project administration, writing - review and editing

Fernando Govantes: Conceptualization, funding acquisition, methodology, project administration, resources, supervision, writing - original draft, writing - review and editing

## REFERENCES

Altegoer F, Schuhmacher J, Pausch P, Bange G. 2014. From molecular evolution to biobricks and synthetic modules: a lesson by the bacterial flagellum. Biotechnol Genet Eng Rev 30:49–64.

Abramson J, Adler J, Dunger J, Evans R, Green T, Pritzel A, Ronneberger O, Willmore L, Ballard AJ, Bambrick J, Bodenstein SW, Evans DA, Hung CC, O’Neill M, Reiman D, Tunyasuvunakool K, Wu Z, Žemgulytė A, Arvaniti E, Beattie C, Bertolli O, Bridgland A, Cherepanov A, Congreve M, Cowen-Rivers AI, Cowie A, Figurnov M, Fuchs FB, Gladman H, Jain R, Khan YA, Low CMR, Perlin K, Potapenko A, Savy P, Singh S, Stecula A, Thillaisundaram A, Tong C, Yakneen S, Zhong ED, Zielinski M, Žídek A, Bapst V, Kohli P, Jaderberg M, Hassabis D, Jumper JM. 2024. Accurate structure prediction of biomolecular interactions with AlphaFold 3. Nature 630):493–500.

Alexander RP, Lowenthal AC, Harshey RM, Ottemann KM. 2010. CheV: CheW-like coupling proteins at the core of the chemotaxis signaling network. Trends Microbiol 18:494–503.

Altschul SF, Gish W, Miller W, Myers EW, Lipman DJ. 1990. Basic local alignment search tool. J Mol Biol 215:403–410.

Alvarado A, Kjær A, Yang W, Mann P, Briegel A, Waldor MK, Ringgaard S. 2017. Coupling chemosensory array formation and localization. Elife 6:e31058.

Bertrand JJ, West JT, Engel JN. 2010. Genetic analysis of the regulation of type IV pilus function by the Chp chemosensory system of *Pseudomonas aeruginosa*. J Bacteriol 192:994–1010.

Briegel A, Ladinsky MS, Oikonomou C, Jones CW, Harris MJ, Fowler DJ, Chang YW, Thompson LK, Armitage JP, Jensen GJ. 2014. Structure of bacterial cytoplasmic chemoreceptor arrays and implications for chemotactic signaling. Elife 3:e02151.

Canarini A, Kaiser C, Merchant A, Richter A, Wanek W. 2019. Root exudation of primary metabolites: mechanisms and their roles in plant responses to environmental stimuli. Front Plant Sci 10:157.

Choi KH, Gaynor JB, White KG, Lopez C, Bosio CM, Karkhoff-Schweizer RR, Schweizer HP. 2005. A Tn7-based broad-range bacterial cloning and expression system. Nat Methods 2:443–448.

Choi KH, Kumar A, Schweizer HP. 2006. A 10-min method for preparation of highly electrocompetent *Pseudomonas aeruginosa* cells: application for DNA fragment transfer between chromosomes and plasmid transformation. J Microbiol Methods 64:391–397.

Colin R, Ni B, Laganenka L, Sourjik V. 2021. Multiple functions of flagellar motility and chemotaxis in bacterial physiology. FEMS Microbiol Rev 45:fuab038.

de Lorenzo V, Pérez-Pantoja D, Nikel PI. 2024. *Pseudomonas putida* KT2440: the long journey of a soil-dweller to become a synthetic biology chassis. J Bacteriol 206(7):e0013624.

Dower WJ, Miller JF, Ragsdale CW. 1988. High efficiency transformation of *E. coli* by high voltage electroporation. Nucleic Acids Res 16:6127–6145.

Ducret A, Quardokus EM, Brun YV. 2016. MicrobeJ, a tool for high throughput bacterial cell detection and quantitative analysis. Nat Microbiol 1:16077.

Espinosa-Urgel M, Salido A, Ramos JL. 2000. Genetic analysis of functions involved in adhesion of *Pseudomonas putida* to seeds. J Bacteriol 182:2363–2369.

García-Fontana C, Reyes-Darias JA, Muñoz-Martínez F, Alfonso C, Morel B, Ramos JL, Krell T. 2013. High specificity in CheR methyltransferase function: CheR2 of *Pseudomonas putida* is essential for chemotaxis, whereas CheR1 is involved in biofilm formation. J Biol Chem 288:18987–18999.

Greenfield D, McEvoy AL, Shroff H, Crooks GE, Wingreen NS, Betzig E, Liphardt J. 2009. Self-organization of the *Escherichia coli* chemotaxis network imaged with super-resolution light microscopy. PLoS Biol 7:e1000137.

Grimm AC, Harwood CS. 1997. Chemotaxis of *Pseudomonas* spp. to the polyaromatic hydrocarbon naphthalene. Appl Environ Microbiol 63:4111–4115.

Hall BA, Armitage JP, Sansom MS. 2012. Mechanism of bacterial signal transduction revealed by molecular dynamics of Tsr dimers and trimers of dimers in lipid vesicles. PLoS Comput Biol 8:e1002685.

Harwood CS, Fosnaugh K, Dispensa M. 1989. Flagellation of *Pseudomonas putida* and analysis of its motile behavior. J Bacteriol 171:4063–4066.

Harwood CS, Rivelli M, Ornston LN. 1984. Aromatic acids are chemoattractants for *Pseudomonas putida*. J Bacteriol 160:622–628.

Hickman JW, Tifrea DF, Harwood CS. 2005. A chemosensory system that regulates biofilm formation through modulation of cyclic diguanylate levels. Proc Natl Acad Sci U S A. 102:14422–7.

Hintsche M, Waljor V, Großmann R, Kühn MJ, Thormann KM, Peruani F, Beta C. 2017. A polar bundle of flagella can drive bacterial swimming by pushing, pulling, or coiling around the cell body. Sci Rep 7:16771.

Inoue H, Nojima H, Okayama H. 1990. High efficiency transformation of *Escherichia coli* with plasmids. Gene 96:23–28.

Jones CW, Armitage JP. 2015. Positioning of bacterial chemoreceptors. Trends Microbiol 23:247–256.

Karimova G, Pidoux J, Ullmann A, Ladant D. 1998. A bacterial two-hybrid system based on a reconstituted signal transduction pathway. Proc Natl Acad Sci U S A 95:5752–5756.

Karimova G, Ullmann A, Ladant D. 2000. A bacterial two-hybrid system that exploits a cAMP signaling cascade in *Escherichia coli*. Methods Enzymol 328:59–73.

Lacal J, Muñoz-Martínez F, Reyes-Darías JA, Duque E, Matilla M, Segura A, Calvo JJ, Jímenez-Sánchez C, Krell T, Ramos JL. 2011. Bacterial chemotaxis towards aromatic hydrocarbons in Pseudomonas. Environ Microbiol 13:1733–1744.

Leal-Morales A, Pulido-Sánchez M, López-Sánchez A, Govantes F. 2022. Transcriptional organization and regulation of the *Pseudomonas putida* flagellar system. Environ Microbiol 24:137–157.

López-Farfán D, Reyes-Darias JA, Matilla MA, Krell T. 2019. Concentration dependent effect of plant root exudates on the chemosensory systems of *Pseudomonas putida* KT2440. Front Microbiol 10:78.

Luu RA, Schneider BJ, Ho CC, Nesteryuk V, Ngwesse SE, Liu X, Parales JV, Ditty JL, Parales RE. 2013. Taxis of *Pseudomonas putida* F1 toward phenylacetic acid is mediated by the energy taxis receptor Aer2. Appl Environ Microbiol 79:2416–2423.

Madeira F, Madhusoodanan N, Lee J, Eusebi A, Niewielska A, Tivey ARN, Lopez R, Butcher S. 2024.The EMBL-EBI Job Dispatcher sequence analysis tools framework in 2024. Nucleic Acids Res 52:521–525.

Mandelbaum RT, Wackett LP, Allan DL.1993. Mineralization of the s-triazine ring of atrazine by stable bacterial mixed cultures. Appl Environ Microbiol 59:1695–701.

Mileykovskaya E, Dowhan W. 2000. Visualization of phospholipid domains in *Escherichia coli* by using the cardiolipin-specific fluorescent dye 10-N-nonyl acridine orange. J Bacteriol 182:1172–1175.

Miller JH. 1993. A short course in bacterial genetics – A laboratory manual and handbook for Escherichia coli and related bacteria. Cold Spring Harbor, New York: Cold Spring Harbor Laboratory Press.

Miyata M, Robinson RC, Uyeda TQP, Fukumori Y, Fukushima SI, Haruta S, Homma M, Inaba K, Ito M, Kaito C, Kato K, Kenri T, Kinosita Y, Kojima S, Minamino T, Mori H, Nakamura S, Nakane D, Nakayama K, Nishiyama M, Shibata S, Shimabukuro K, Tamakoshi M, Taoka A, Tashiro Y, Tulum I, Wada H, Wakabayashi KI. 2020. Tree of motility - A proposed history of motility systems in the tree of life. Genes Cells 25:6–21.

Muok AR, Briegel A, Crane BR. 2020. Regulation of the chemotaxis histidine kinase CheA: a structural perspective. Biochim Biophys Acta Biomembr 1862:183030.

Navarrete B, Leal-Morales A, Serrano-Ron L, Sarrió M, Jiménez-Fernández A, Jiménez-Díaz L, López-Sánchez A, Govantes F. 2019. Transcriptional organization, regulation and functional analysis of *flhF* and *fleN* in *Pseudomonas putida*. PLoS One 14:e0214166.

Nie H, Xiao Y, Song M, Wu N, Peng Q, Duan W, Chen W, Huang Q. 2022. Wsp system oppositely modulates antibacterial activity and biofilm formation via FleQ-FleN complex in *Pseudomonas putida*. Environ Microbiol 24:1543–1559.

Osterberg S, Skärfstad E, Shingler V. 2010. The sigma-factor FliA, ppGpp and DksA coordinate transcriptional control of the *aer2* gene of *Pseudomonas putida*. Environ Microbiol 12:1439–1451.

Parales RE, Ditty JL. 2018. Chemotaxis to atypical chemoattractants in soil bacteria. Methods Mol Biol 1729:255–280.

Parkinson JS. 1976. *cheA*, *cheB*, and *cheC* genes of *Escherichia coli* and their role in chemotaxis. J Bacteriol 126:758–770.

Pulido-Sánchez M, Leal-Morales A, López-Sánchez A, Cava F, Govantes F. 2025. Spatial, temporal and numerical regulation of polar flagella assembly in *Pseudomonas putida*. Microbiol Res 292:128033.

Rico-Jiménez M, Roca A, Krell T, Matilla MA. 2022. A bacterial chemoreceptor that mediates chemotaxis to two different plant hormones. Environ Microbiol 24:3580–3597.

Ringgaard S, Schirner K, Davis BM, Waldor MK. 2011. A family of ParA-like ATPases promotes cell pole maturation by facilitating polar localization of chemotaxis proteins. Genes Dev 25:1544–1555.

Ringgaard S, Yang W, Alvarado A, Schirner K, Briegel A. 2018. Chemotaxis arrays in *Vibrio* species and their intracellular positioning by the ParC/ParP system. J Bacteriol 200:e00793–17.

Ringgaard S, Zepeda-Rivera M, Wu X, Schirner K, Davis BM, Waldor MK. 2014. ParP prevents dissociation of CheA from chemotactic signaling arrays and tethers them to a polar anchor. Proc Natl Acad Sci U S A 111:255–264.

Rodríguez-Herva JJ, Duque E, Molina-Henares MA, Navarro-Avilés G, Van Dillewijn P, De La Torre J, Molina-Henares AJ, La Campa AS, Ran FA, Segura A, Shingler V, Ramos JL. 2010. Physiological and transcriptomic characterization of a *fliA* mutant of *Pseudomonas putida* KT2440. Environ Microbiol Rep 2:373–380.

Rossmann F, Brenzinger S, Knauer C, Dörrich AK, Bubendorfer S, Ruppert U, Bange G, Thormann KM. 2015.The role of FlhF and HubP as polar landmark proteins in *Shewanella putrefaciens* CN-32. Mol Microbiol 98:727–742.

Rudner DZ, Losick R. 2010. Protein subcellular localization in bacteria. Cold Spring Harb Perspect Biol 2:a000307.

Sambrook JE, Russell DW. 2000. Molecular cloning, a laboratory manual. Cold Spring Harbor, New York, USA: Cold Spring Harbor Laboratory Press.

Sarand I, Osterberg S, Holmqvist S, Holmfeldt P, Skärfstad E, Parales RE, Shingler V. 2008. Metabolism-dependent taxis towards (methyl)phenols is coupled through the most abundant of three polar localized Aer-like proteins of *Pseudomonas putida*. Environ Microbiol 10:1320–1334.

Scheller C, Krebs F, Wiesner R, Wätzig H, Oltmann-Norden I. 2021. A comparative study of CE-SDS, SDS-PAGE, and Simple Western-Precision, repeatability, and apparent molecular mass shifts by glycosylation. Electrophoresis 42:1521–1531.

Schmidt LM, Pulido-Sánchez M, Treuner-Lange A, Zehner L, López-Sánchez A, Cava F, Govantes F, Thormann KM. 2026. Functional characterization of the polar organizer protein FimV in *Pseudomonas putida*. J Bacteriol 208:e0049725.

Schneider CA, Rasband WS, Eliceiri KW. 2012. NIH Image to ImageJ: 25 years of image analysis. Nat Methods 9:671–675.

Schramm A, Bignon C, Brocca S, Grandori R, Santambrogio C, Longhi S. 2019. An arsenal of methods for the experimental characterization of intrinsically disordered proteins - How to choose and combine them? Arch Biochem Biophys. 676:108055.

Shiomi D, Yoshimoto M, Homma M, Kawagishi I. 2006. Helical distribution of the bacterial chemoreceptor via colocalization with the Sec protein translocation machinery. Mol Microbiol 60:894–906.

Sourjik V, Wingreen NS. 2012. Responding to chemical gradients: bacterial chemotaxis. Curr Opin Cell Biol 24:262–268.

Strahl H, Ronneau S, González BS, Klutsch D, Schaffner-Barbero C, Hamoen LW. 2015. Transmembrane protein sorting driven by membrane curvature. Nat Commun 6:8728.

Surovtsev IV, Jacobs-Wagner C. 2018. Subcellular organization: a critical feature of bacterial cell replication. Cell 172:1271–1293.

Tiwari P, Kaila P, Guptasarma P. Understanding anomalous mobility of proteins on SDS-PAGE with special reference to the highly acidic extracellular domains of human E- and N-cadherins. 2019. Electrophoresis 40:1273–1281.

Thiem S, Kentner D, Sourjik V. 2007. Positioning of chemosensory clusters in *E. coli* and its relation to cell division. EMBO J 26:1615–23.

Thiem S, Sourjik V. 2008. Stochastic assembly of chemoreceptor clusters in *Escherichia coli*. Mol Microbiol 68:1228–1236.

Thompson SR, Wadhams GH, Armitage JP. 2006. The positioning of cytoplasmic protein clusters in bacteria. Proc Natl Acad Sci U S A 103:8209–8214.

Wadhams GH, Armitage JP. 2004. Making sense of it all: bacterial chemotaxis. Nat Rev Mol Cell Biol 5:1024–1037.

Waterhouse AM, Procter JB, Martin DMA, Clamp M, Barton GJ. 2009. Jalview Version 2-a multiple sequence alignment editor and analysis workbench. Bioinformatics 25: 1189–1191.

Whitchurch CB, Leech AJ, Young MD, Kennedy D, Sargent JL, Bertrand JJ, Semmler AB, Mellick AS, Martin PR, Alm RA, Hobbs M, Beatson SA, Huang B, Nguyen L, Commolli JC, Engel JN, Darzins A, Mattick JS. 2004. Characterization of a complex chemosensory signal transduction system which controls twitching motility in *Pseudomonas aeruginosa*. Mol Microbiol 52:873–93.

Yamaichi Y, Bruckner R, Ringgaard S, Möll A, Cameron DE, Briegel A, Jensen GJ, Davis BM, Waldor MK. 2012. A multidomain hub anchors the chromosome segregation and chemotactic machinery to the bacterial pole. Genes Dev 26:2348–60.

Yang W, Briegel A. 2020. Diversity of bacterial chemosensory arrays. Trends Microbiol 28:68–80.

