## Supplemental Materials for "FimV, ParC and ParP coordinate polar assembly of the chemosensory arrays in *Pseudomonas putida*"

**_____________________________________________________________**

**SUPPLEMENTARY INFORMATION**

**1. SUPPLEMENTARY MATERIALS AND METHODS**

**Plasmid and strain construction**

**Construction of MRB113, MRB213, MRB225, MRB228, MRB236 and MRB238.** To construct *P. putida* in-frame deletion mutants of the *parC*, *parP* and *cheA* ORFs, upstream and downstream chromosomal regions flanking these genes were PCR-amplified using appropriate oligonucleotide pairs specified in **Table S1**. Upstream and downstream *parC* PCR products were cleaved with EcoRI and BamHI (upstream region) or BamHI and XbaI (downstream region) and three-way ligated into EcoRI- and XbaI-digested gene replacement vector pEMG that incorporates two I-SceI sites flanking the *lacZ* polylinker, yielding pMRB288. Upstream and downstream *parP* PCR products were cleaved with XmaI and BamHI (upstream region) or BamHI and XbaI (downstream region), and three-way ligated into XmaI- and XbaI-digested pEMG, yielding pMRB447. Upstream and downstream *cheA* PCR products were cleaved with EcoRI and HindIII (upstream region) or HindIII and XbaI (downstream region), and three-way ligated into EcoRI- and XbaI-digested pEMG, yielding pMRB355. These vectors were introduced by triparental mating into *P. putida* KT24442 harbouring the pSW-I plasmid, expressing the I-SceI endonuclease from the 3MB-inducible *xylS*-P*m* system. Selection of integration and allelic exchange by homologous recombination-mediated repair of the chromosomal cleavage at the integrated I-SceI site was conducted as previously described **(Martínez-García and de Lorenzo, 2011)** to generate ∆*parC* (MRB113)*,* ∆*parP* (MRB236) and ∆*cheA* (MRB213) mutant strains, respectively. Similarly, pMRB355 was transferred to ∆*parC,* ∆*parP* and ∆*fimV* (MRB97) to generate ∆*parC*∆*cheA* (MRB228), ∆*parP*∆*cheA* (MRB238) and ∆*fimV*∆*cheA* (MRB225) double mutants. Loci deletions were verified by PCR. Curation of pSW-I plasmid was carried out by successive growth cycles in LB medium.

**Construction of pMRB450 and pMRB473.** For ParC-GFP and ParP-GFP expression under the P*sal* promoter, pMRB189-derivatives were constructed by cloning the PCR-amplified *parC* and *parP* ORFs into BamHI- and XbaI-digested, or BamHI- and XmaI-digested pMRB189, yielding pMRB473 and pMRB450, respectively.

**Construction of pMRB481.** For CheA-GFP expression under its natural P*cheA* promoter, a 2780 bp P*cheA-cheA* chromosomal region genomic region of *P. putida* KT2440 (GenBank: NC_002947.4, 4.927.931-4.930.710) flanked by SpeI and BamHI restriction sites was commercially synthesized (NZYTech). The resulting DNA fragment was cloned into SpeI- and BamHI-digested pMRB187, yielding pMRB481.

**Construction of pMRB457, pMRB461, pMRB462, pMRB464, pMRB468 and pMRB469.** For T18- and T25-hybrid proteins expression for BACTH assays, pUT18- and pKNT25-derivatives were constructed by cloning the PCR-amplified *fimV*, *parC* and *parP* ORFs into BamHI- and HindIII-digested pUT18 or pKNT25, yielding pMRB457 (FimV-T18), pMRB461 (ParC-T18), pMRB462 (ParP-T18), pMRB464 (FimV-T25), pMRB468 (ParC-T25) and pMRB469 (ParP-T25).

**Genomic integration of pVI812.** For Aer2-YFP expression, suicide plasmid pVI812 carrying *aer2-yfp,* was integrated into the genome of *P. putida* KT2442 wild-type and mutant strains by homologous recombination as described above. Single-site recombinant was maintained relying on kanamycin selection as previously described **(Sarand *et al*., 2008)**.

**2. SUPPLEMENTARY TABLE**

**Table S1. Bacterial strains, plasmids and oligonucleotides used in this work.** Cm: chloramphenicol. Rif: rifampicin. Km: kanamycin. Ap: ampicillin. Gm: gentamycin. Sm: streptomycin.

**Bacterial strain	Genotype/phenotype	Reference/source**

***E. coli***

BTH101	F-, *cya-99*, *araD139, galE15, galK16, rpsL1* (Sm^r^), *hsdR2, mcrA1, mcrB1*	Karimova *et al*., 2000

DH5α	Φ80d*lacZ*∆M15 ∆(*lacZYA-argF*)U169 *recA*1 *endA*1 *hsdR*17 (r_k_^-^ m_k_^+^) *supE*44 *thi*-1 *gyrA* *relA*1	Hanahan, 1983

DH5α λpir	DH5α with lysogenic phage λ-pir, host for R6K replication origin plasmids	Víctor de Lorenzo

***P. putida***

KT2442	mt-2 *hsdR*1 (r^-^ m^+^). Cm^r^ Rif^r^.	Franklin *et al*., 1981

MRB113	KT2442 ∆*parC*. Cm^r^ Rif^r^.	This work

MRB213	KT2442 ∆*cheA*. Cm^r^ Rif^r^.	This work

MRB225	KT2442 ∆*fimV*∆*cheA*. Cm^r^ Rif^r^	This work

MRB228	KT2442 ∆*parC*∆*cheA*. Cm^r^ Rif^r^	This work

MRB236	KT2442 ∆*parP*. Cm^r^ Rif^r^	This work

MRB238	KT2442 ∆*parP*∆*cheA*. Cm^r^ Rif^r^	This work

MRB52	KT2442 ∆*fleQ*. Cm^r^ Rif^r^.	Navarrete *et al*., 2019

MRB69	KT2442 ∆*flhF*. Cm^r^ Rif^r^.	Navarrete *et al*., 2019

MRB71	KT2442 ∆*fleN*. Cm^r^ Rif^r^.	Navarrete *et al*., 2019

MRB97	KT2442 ∆*fimV*. Cm^r^ Rif^r^.	Pulido-Sánchez *et al*., 2025

**Plasmid	Genotype/phenotype	Reference/source**

pEMG	pJP5603 bearing a *lacZ*α polylinker with two flanking I-SceI sites. R6K, Mob^+^, Km^r^	Martínez-García and de Lorenzo, 2011

pKNT25	BACTH expression vector for T25 fragment C-terminal translational fusions under the P*lac* promoter. p15A, Km^r^	Karimova *et al*., 2000

pKT25	BACTH expression vector for T25 fragment N-terminal translational fusions under the P*lac* promoter. p15A, Km^r^	Karimova *et al*., 2000

pKT25-zip	pKT25-derived vector containing a leucine zipper domain, positive control in BACTH assays. Km^r^	Karimova *et al*., 2000

pMRB172	pUC18Sfi-miniTn*7*BB-Gm-based delivery plasmid for miniTn*7*BB-Gm [*nahR*-P*sal*]. Ap^r^ Gm^r^	Leal-Morales *et al*., 2022

pMRB187	pUC18Sfi-miniTn*7*BB-Gm-based delivery plasmid for C-terminal *gfp-*mut3 translational fusions. Ap^r^ Gm^r^	Pulido-Sánchez *et al.,* 2025

pMRB189	pMRB172-derived delivery plasmid for C-terminal *gfp-*mut3 translational fusions. Ap^r^ Gm^r^	Pulido-Sánchez *et al.,* 2025

pMRB200	pMRB187-derived vector containing P*flhF*-*flhF* translationally fused to *gfp*mut3 at its C-terminus. Ap^r^ Gm^r^	Pulido-Sánchez *et al.,* 2025

pMRB236	pMRB189-derived vector containing a RBS site and *fimV* translationally fused to *gfp*mut3 at its C-terminus. Ap^r^ Gm^r^	Pulido-Sánchez *et al.,* 2025

pMRB238	pMRB172-derived vector containing a RBS site and *gfp*mut3 N-terminal translational fusions Ap^r^ Gm^r^.	Pulido-Sánchez *et al.,* 2025

pMRB288	pEMG bearing a 1800 bp insert with upstream and downstream flanking regions of *parC*. Km^r^	This work

pMRB302	pUC18Sfi-mini*Tn*7BB-Gm-derived vector containing P*fliC*-*fliC^S267C^.* Ap^r^ Gm^r^	Pulido-Sánchez *et al.,* 2025

pMRB355	pEMG bearing a 1033 bp insert with upstream and downstream flanking regions of *cheA*. Km^r^	This work

pMRB375	pMRB238-derived vector containing *fleN* translationally fused to *gfp*mut3 at its N-terminus. Ap^r^ Gm^r^	Pulido-Sánchez *et al.,* 2025

pMRB447	pEMG bearing a 1059 bp insert with upstream and downstream flanking regions of *parP*. Km^r^	This work

pMRB450	pMRB189-derived vector containing a RBS site and *parP* translationally fused to *gfp*mut3 at its C-terminus. Ap^r^ Gm^r^This work

pMRB457	pUT18-derived vector containing *fimV* translationally fused to T18 at its C-terminus. Ap^r^	This work

pMRB461	pUT18-derived vector containing *parC* translationally fused to T18 at its C-terminus. Ap^r^	This work

pMRB462	pUT18-derived vector containing *parP* translationally fused to T18 at its C-terminus. Ap^r^	This work

pMRB464	pKNT25-derived vector containing *fimV* translationally fused to T25 at its C-terminus. Km^r^	This work

pMRB468	pKNT25-derived vector containing *parC* translationally fused to T25 at its C-terminus. Km^r^	This work

pMRB469	pKNT25-derived vector containing *parP* translationally fused to T25 at its C-terminus. Km^r^	This work

pMRB473	pMRB189-derived vector containing a RBS site and *parC* translationally fused to *gfp*mut3 at its C-terminus. Ap^r^ Gm^r^This work

pMRB481	pMRB187-derived vector containing P*cheA-cheA* translationally fused to *gfp*mut3 at its C-terminus. Ap^r^ Gm^r^	This work

pRK2013	Helper plasmid for triparental mating. ColE1, Mob^+^, Km^r^	Figurski and Helinski, 1979

pSW-I	Plasmid expressing I-SceI from *xylS*-P*m*. RK2, Mob^+^, Ap^r^	Wong and Mekalanos, 2000

pTNS2	Helper plasmid expressing the Tn7 transposase. R6K, Mob^+^, Ap^r^	Choi *et al*., 2005

pUC18Sfi-miniTn*7*BB-Gm	pUC18SfiI-based delivery plasmid for the synthetic minitransposon mini*Tn*7BB-Gm. Ap^r^ Gm^r^	Jiménez-Fernández *et al*., 2014

pUT18	BACTH expression vector for T18 fragment C-terminal translational fusions under the P*lac* promoter. ColE1, Ap^r^	Karimova *et al*., 2000

pUT18-zip	pUT18-derived vector containing a leucine zipper domain, positive control in BACTH assays. Ap^r^	Karimova *et al*., 2000

pVI812	3′ *aer2–yfp* on pDM4-Km. Km^r^	Sarand *et al*., 2008

**Oligonucleotide	Sequence (5’ to 3’)	Use	____**

CheA -DOWN-DOWN_rev	GTTTGTAGGGTTTGCGCT	*cheA* chromosomal deletion verification

CheA -DOWN-HindIII_fwd	GATCAAGCTTTGATTTCGGTGGCGCG	*cheA* downstream region amplification

CheA -DOWN-XbaI_rev	GATCTCTAGAGCCTGGCTGGTAAAGGTGC	*cheA* downstream region amplification

CheA -UP-EcoRI_fwd	GATCGAATTCCACCATGGACCTGGTCG	*cheA* upstream region amplification

CheA-UP-HindIII_rev	GATCAAGCTTCATCAAACGTGCTCCTTAAA	*cheA* upstream region amplification

CheA -UP-UP_fwd	GTGGTCAAGCTCACCGAG	*cheA* chromosomal deletion verification

FimV-BACTH-BamHI_rev	CAGTGGATCCCCGACCAGCCGGGAGAGCAT	*fimV* ORF amplification for BACTH

FimV-BACTH-HindIII_fwd	CAGTAAGCTTACTTCGAATTCGCAAACTGGTTC	*fimV* ORF amplification for BACTH

ParC-BamHI_rev	CAGTGGATCCGGCTACCTGCACTGCGTTC	*parC* ORF amplification

ParC-BACTH-BamHI_rev	CAGTGGATCCCCGGCTACCTGCACTGCGTT	*parC* ORF amplification for BACTH

ParC-BACTH-HindIII_fwd	CAGTAAGCTTACGCGAGCTTTCGAATTTTG	*parC* ORF amplification for BACTH

ParC-DOWN-BamHI_fwd	ACGTGGATCCGCTGCTCAAGCACCTGCT	*parC* downstream region amplification

ParC-DOWN-DOWN_rev	TTTGGCGTGCATCTGCTT	*parC* chromosomal deletion verification

ParC-DOWN-XbaI_rev	ACGTTCTAGAGACGACTTTTTCATTCCCCTACC	*parC* downstream region amplification

ParC-UP-BamHI_rev	ACGTGGATCCCAAGCAAACTCTATCTATTGATTTGGC	*parC* upstream region amplification

ParC-UP-EcoRI-KpnI_fwd	ACGTGAATCCGGTACCGCTGGTCAAAGACAGCGAGC	*parC* upstream region amplification

ParC-UP-UP_fwd	ATTGGCGATGAGCAGCC	*parC* chromosomal deletion verification

ParC-XbaI-RBS_fwd	CAGTTCTAGAGAAAGAGGAGAAATACTAGTTGCGCGAGCTTTCGAAT	*parC* ORF amplification

ParP-BACTH-BamHI_rev	CAGTGGATCCCCTTTCTGTTTGGCGTGCATCT	*parP* ORF amplification for BACTH

ParP-BACTH-HindIII_fwd	CAGTAAGCTTAACTCAAACCCGGCAAACC	*parP* ORF amplification for BACTH

ParP-BamHI_rev	CAGTGGATCCTTTCTGTTTGGCGTGCATCTG	*parP* ORF amplification

ParP-DOWN-BamHI_fwd	GATCGGATCCTGACACAGAACATACCGCCAAC	*parP* downstream region amplification

ParP-DOWN-DOWN_rev	CCAGCGCCAGGAAGATGAC	*parP* chromosomal deletion verification

ParP-DOWN-XbaI_rev	CATGTCTAGAGGACCACTCTTCCTCGGTCA	*parP* downstream region amplification

ParP-UP-BamHI_rev	CATGGGATCCCATCAGGCTACCTGCACTGC	*parP* upstream region amplification

ParP-UP-XmaI_fwd	CATGCCCGGGGAGCGCATTTCGCTGTTG	*parP* upstream region amplification

ParP-UP-UP_fwd	ACTGCTGCTGCCCACCAG	*parP* chromosomal deletion verification

ParP-XmaI-RBS_fwd	CAGTCCCGGGGAAAGAGGAGAAATACTAGATGACTCAAACCCGGCAAACC	*parP* ORF amplification

Tn7-GlmS	AATCTGGCCAAGTCGGTGAC	Confirmation of miniTn7-derivatives integration

Tn7-R109	CAGCATAACTGGACTGATTTCAG	Confirmation of miniTn7-derivatives integration

**3. SUPPLEMENTARY FIGURES**

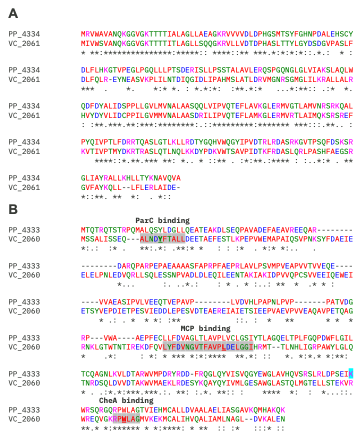

**Figure S1. Sequence alignments of ParC and ParP in *P. putida* and *V. cholerae*. A.** Alignment of ParC in *P. putida* (PP_4334) and *V. cholerae* (VC_2061). **B.** Alignment of ParP in *P. putida* (PP_4333) and *V. cholerae* (VC_2060). The proposed ParC-, MCP- and CheA-binding motifs are shaded, and the residues identified as essential are in bold and underlined.

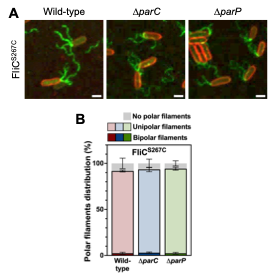

**Figure S2. Flagellar filaments in the ∆*parC* and ∆*parP* mutants. A.** Confocal microscopy images of wild-type, ∆*parC* and ∆*parP* cells expressing P*fliC-fliC^S267C^* (green). Alexa Fluor^TM^ 488 C_5_ maleimide was used for FliC^S267C^ filament staining. FM^TM^ 4-64 was used as membrane stain (red). Images are shown as the maxima projections of seven Z-sections of the yellow channel, merged with the red channel showing the cell contour at the focal plane. Scale bar: 2 µm. **B.** Frequency of cells bearing bipolar, unipolar o no polar FliC^S267C^ filaments. Columns and error bars represent the averages and standard deviations of at least three separate replicates (n = 500 cells). Welch's *t*-test did not detect significant differences (*p*≥0.05).

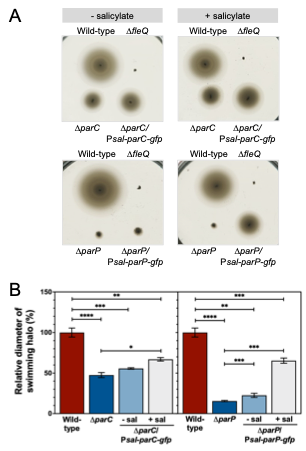

**Figure S3. Complementation of the ∆*parC* and ∆*parP* mutants. A.** Soft agar-based swimming motility assays showing complementation of the ∆*parC* and ∆*parP* mutants with P*sal-parC-gfp* and P*sal-parP-gfp*, respectively, in the absence (left) or in the presence (right) of 2 mM salicylate. The wild-type strain and the ∆*fleQ* mutant were assayed in the same plates as positive and negative controls. Each picture shows a representative swim plate out of at least three biological replicates. **C.** Relative diameters of swimming halos. Values are normalized to the wild-type (100%). Columns and error bars represent the averages and standard deviations of at least three biological replicates. Stars designate for *p*-values for Welch's *t*-test (*:*p*<0.05; **:*p*<0.01; ***:*p*<0.001; ****:*p*<0.0001).

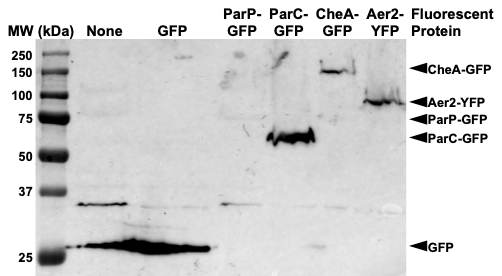

**Figure S4. Western blot of KT2442 producing fluorescent fusion proteins**. Extracts from KT2442 producing no fluorescent protein, GFP, or the ParP-GFP, ParC-GFP, CheA-GFP or Aer2-YFP fusion proteins were resolved on SDS-PAGE, blotted and probed with anti-GFP antiserum. Arrowheads indicate the positions of the fluorescent proteins detected.

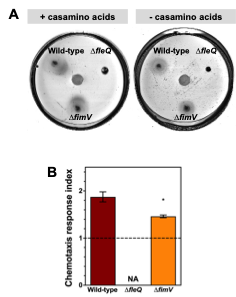

**Figure S5. Chemotaxis assay of the ∆*fimV* mutant. A.** Chemical-in-plug chemotaxis assays of the ∆*fimV* mutant towards an agarose plug with (left) or without (right) 0.2% casamino acids at the center of the plate. The wild-type strain and the ∆*fleQ* mutant were assayed in the same plates as positive and negative controls. Each picture shows a representative swim plate out of at least three biological replicates. **B.** Chemotaxis response indexes of the aforementioned strains when exposed to an agarose plug containing casamino acids. Columns and error bars represent the averages and standard deviations of at least three biological replicates. NA: not applicable. Stars designate *p*-values for Welch's *t*-test (*:*p*<0.05).

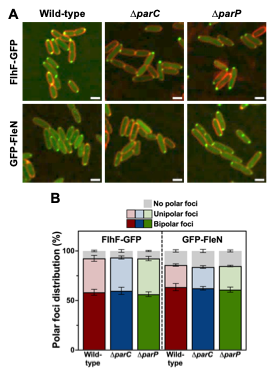

**Figure S6. Intracellular location of FlhF and FleN in the ∆*parC* and ∆*parP* mutants. A.** Confocal microscopy images of wild-type, ∆*parC* and ∆*parP* cells expressing P*sal-flhF-gfp* or P*sal-gfp-fleN* (green) in the presence of 2 mM salicylate. FM^TM^ 4-64 was used as membrane stain (red). Images are shown as the maxima projections of seven Z-sections of the green channel, merged with the red channel showing the cell contour at the focal plane. Scale bar: 2 µm. **B.** Frequency of cells of the aforementioned strains bearing bipolar, unipolar o no polar FlhF-GFP or GFP-FleN foci. Columns and error bars represent the averages and standard deviations of at least three separate replicates (n = 500 cells). Welch's *t*-test did not detect significant differences (*p*≥0.05).

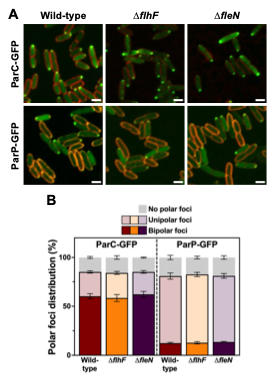

**Figure S7. Intracellular location of ParC and ParP in the ∆*flhF* and ∆*fleN* mutants. A.** Confocal microscopy images of wild-type, ∆*flhF* and ∆*fleN* cells expressing P*sal-parC-gfp* or P*sal-parP-gfp* (green) in the presence of 2 mM salicylate. FM^TM^ 4-64 was used as membrane stain (red). Images are shown as the maxima projections of seven Z-sections of the green channel, merged with the red channel showing the cell contour at the focal plane. Scale bar: 2 µm. **B.** Frequency of cells of the aforementioned strains bearing bipolar, unipolar o no polar ParC-GFP or ParP-GFP foci. Columns and error bars represent the averages and standard deviations of at least three separate replicates (n = 500 cells). Welch's *t*-test did not detect significant differences (*p*≥0.05).

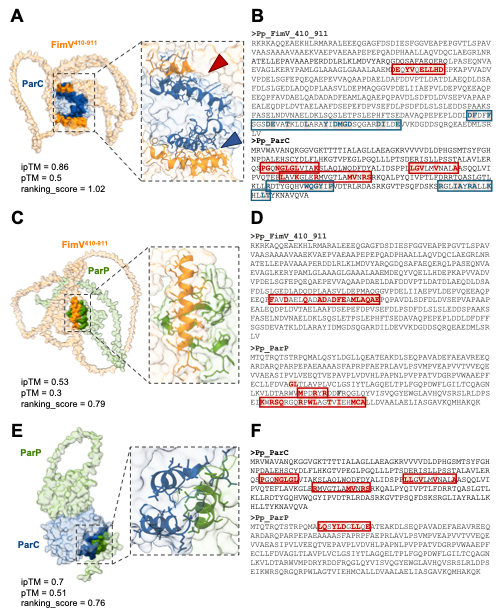

**Figure S8. AlphaFold3 models of FimV, ParC and ParP interactions.** **A, C and E.** AlphaFold3-generated models of the interactions between FimV^410-911^ and ParC **(A)**, FimV^410-911^ and ParP **(C)**, or ParC and ParP **(E)**. Images represent the surface areas of the interacting proteins. Insets show a detail view of the interaction surfaces with the contact regions highlighted as secondary structure cartoons with side chains. A single model with the highest quality scores (shown next to each image as a reference) out of five analyzed is shown. **B, D and F.** Sequences of the interacting protein pairs FimV^410-911^ and ParC **(B)**, FimV^410-911^ and ParP **(D)**, or ParC and ParP **(F)**, showing the interacting regions and residues. Consensus contacts, present in at least four out of five models evaluated are shown in bold and highlighted. Boxes denote interaction regions containing multiple contacts. Color-coded arrowheads in **A.** and boxes in **B.** denote two separate interaction surfaces. The complete dataset of the structures analyzed is provided as **Supplementary Data 1**.

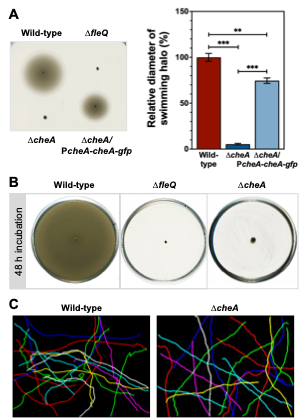

**Figure S9. Swimming motility of the ∆*cheA* mutant. A.** Soft agar-based swimming motility assay showing complementation of the ∆*cheA* mutant with P*cheA-cheA-gfp*. Pictures of representative swim plates (left) and relative diameters of swimming halos (right). Values are normalized to the wild-type (100%). Columns and error bars represent the averages and standard deviations of at least three biological replicates. Stars designate for *p*-values for Welch's *t*-test (**:*p*<0.01; ***:*p*<0.001). **B.** Soft agar-based swimming motility assay of the ∆*cheA* mutant after 48 h incubation. The wild-type strain and the ∆*fleQ* mutant were assayed as positive and negative controls in panels **A** and **B**. **C.** Near-surface swimming trajectories of wild-type and ∆*cheA* cells. Each coloured line represents the trajectory of a single cell tracked for 15 s from brightfield microscopy videos (n = 20 cells).

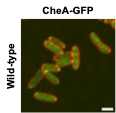

**Figure S10. Intracellular location of CheA in the wild-type strain.** Confocal microscopy images of wild-type cells expressing P*cheA-cheA-gfp* (green). FM^TM^ 4-64 was used as membrane stain (red). Images are shown as the maxima projections of seven Z-sections of the green channel, merged with the red channel showing the cell contour at the focal plane. Scale bar: 2 µm.

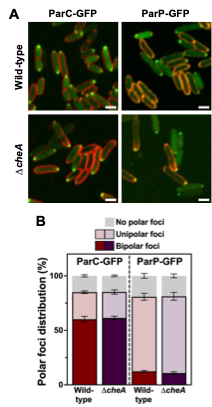

**Figure S11. Intracellular location of ParC and ParP in the ∆*cheA* mutant. A.** Confocal microscopy images of wild-type and ∆*cheA* cells expressing P*sal-parC-gfp* or P*sal-parP-gfp* (green) in the presence of 2 mM salicylate. FM^TM^ 4-64 was used as membrane stain (red). Images are shown as the maxima projections of seven Z-sections of the green channel, merged with the red channel showing the cell contour at the focal plane. Scale bar: 2 µm. **B.** Frequency of cells of the aforementioned strains bearing bipolar, unipolar o no polar ParC-GFP or ParP-GFP foci. Columns and error bars represent the averages and standard deviations of at least three separate replicates (n = 500 cells). Welch's *t*-test did not detect significant differences (*p*≥0.05).

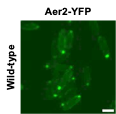

**Figure S12. Intracellular location of MCPs without membrane stain.** Confocal microscopy images of wild-type cells expressing *aer2-yfp* (green). Images are shown as the maxima projections of seven Z-sections of the yellow channel. Scale bar: 2 µm.

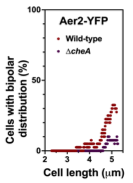

**Figure S13. Cell cycle-dependent bipolar distribution of Aer2.** Frequency of cells of the wild-type strain and the ∆*cheA* mutant displaying bipolar distribution of Aer2-YFP foci *vs.* cell length (n = 500 cells). The plot represents frequency of bipolar distribution in overlapping sets of 40 length-sorted cells plotted against the average length of each set (n = 500 cells).

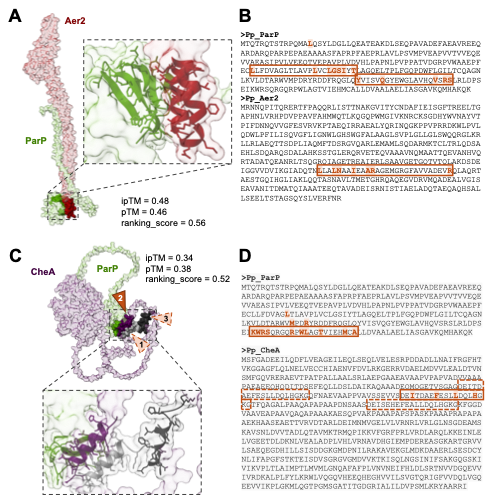

**Figure S14. AlphaFold3 models of ParP interactions with Aer2 and CheA.** **A and C.** AlphaFold3-generated models of the interactions between ParP and Aer2 **(A)**, or ParP and CheA **(C)**. Images represent the surface areas of the interacting proteins. Insets show a detail view of the interaction surfaces with the contact regions highlighted as secondary structure cartoons with side chains. A single model with the highest quality scores (shown next to each image as a reference) out of five analyzed is shown. **B and D.** Sequences of the interacting protein pairs ParP and Aer2 **(B)**, or ParP and CheA **(D)**, showing the interacting regions and residues. Consensus contacts, present in at least four out of five models evaluated are shown in bold and highlighted. Closed boxes denote interaction regions containing multiple contacts. Dashed arrowheads in **C.** and dashed boxes in **D.** denote two additional repeat regions shown to act as alternative contact regions. The complete dataset of the structures analyzed is provided as **Supplementary Data 1**.

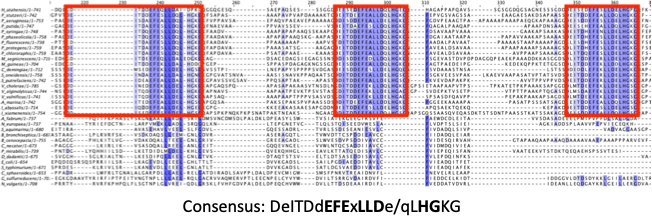

**Figure S15. Alignment of the putative ParP-binding motifs in CheA.** Sequences surrounding the putative ParP-binding repeats (red boxes) in thirty-three bacterial species also containing (top 20) or not containing (bottom 13) ParC and ParP orthologs.

**4. SUPPLEMENTARY REFERENCES**

**1.** Choi KH, Gaynor JB, White KG, Lopez C, Bosio CM, Karkhoff-Schweizer RR, Schweizer HP. 2005. A Tn*7*-based broad-range bacterial cloning and expression system. Nat Methods 2(6):443-448.

**2.** Figurski DH, Helinski DR. 1979. Replication of an origin-containing derivative of plasmid RK2 dependent on a plasmid function provided in *trans*. PNAS 76(4): 1648-1652.

**3.** Franklin FC, Bagdasarian M, Bagdasarian MM, Timmis KN. 1981. Molecular and functional analysis of the TOL plasmid pWWO from *Pseudomonas* *putida* and cloning of genes for the entire regulated aromatic ring meta cleavage pathway. PNAS 78(12):7458-7462.

**4.** Hanahan D. 1983. Studies on transformation of *Escherichia* *coli* with plasmids. J Mol Biol 166(4):557-580.

**5.** Jiménez-Fernández A, López-Sánchez A, Calero P, Govantes F. 2014. The c-di-GMP phosphodiesterase BifA regulates biofilm development in *Pseudomonas* *putida*. Environ Microbiol Rep 7(1):78-84.

**6.** Karimova G, Ullmann A, Ladant D. 2000. A bacterial two-hybrid system that exploits a cAMP signaling cascade in *Escherichia* *coli*. *In* Methods in Enzymology vol 328 p 59-73. Elsevier.

**7.** Leal-Morales A, Pulido-Sánchez M, López-Sánchez A, Govantes F. 2022. Transcriptional organization and regulation of the *Pseudomonas* *putida* flagellar system. Environ Microbiol 24(1):137-157.

**8.** Martínez-García E, De Lorenzo V. 2011. Engineering multiple genomic deletions in Gram-negative bacteria: Analysis of the multi-resistant antibiotic profile of *Pseudomonas* *putida* KT2440. Environ Microbiol 13(10):2702-2716.

**9.** Navarrete B, Leal-Morales A, Serrano-Ron L, Sarrió M, Jiménez-Fernández A, Jiménez-Díaz L, López-Sánchez A, Govantes F. 2019. Transcriptional organization, regulation and functional analysis of *flhF* and *fleN* in *Pseudomonas* *putida*. PLOS ONE 14(3):e0214166.

**10.** Pulido-Sánchez M, Leal-Morales A, López-Sánchez A, Cava F, Govantes F. 2025. Spatial, temporal and numerical regulation of polar flagella assembly in *Pseudomonas* *putida*. Microbiol Res 292:128033.

**11.** Sarand I, Österberg S, Holmqvist S, Holmfeldt P, Skärfstad E, Parales RE, Shingler V. 2008. Metabolism-dependent taxis towards (methyl)phenols is coupled through the most abundant of three polar localized Aer-like proteins of *Pseudomonas putida*. Environ Microbiol 10(5):1320-1334.

**12.** Wong SM, Mekalanos JJ. 2000. Genetic footprinting with mariner-based transposition in *Pseudomonas* *aeruginosa*. PNAS 97(18):10191-10196.
